# Using aerial thermography to map terrestrial thermal environments in unprecedented detail

**DOI:** 10.1101/2025.02.21.638895

**Authors:** Karla Alujević, Guillermo Garcia-Costoya, Noa Ratia, Emma Schmitz, Russell S. Godkin, Akhila C. Gopal, Jelena Bujan, Michael L. Logan

**Author notes:** **Corresponding author:** Karla Alujević. Authors contributed equally (Karla Alujević, Guillermo Garcia-Costoya).

## Abstract

1. The accurate quantification of thermal environments is crucial for predicting the impacts of climate change across ecosystems.
2. Two major obstacles exist in mapping biologically relevant thermal landscapes: 1) overcoming the mismatch between the spatial resolution at which environmental data are typically collected and the scale at which a particular organism experiences thermal variation, and 2) quantifying thermal landscapes without substantial measurement gaps in time or space.
3. We present a new method that integrates aerial thermography from uncrewed aerial vehicles with field-deployed operative temperature models to generate fine-scale, spatiotemporally complete maps of operative temperature.
4. To ensure the accessibility of our method, we developed an R package, ‘`throne`’, which streamlines the necessary corrections to raw drone data and produces operative thermal landscapes for any day or time during which data loggers were deployed.
5. Our method allows researchers to generate detailed and biologically relevant thermal landscapes for species of interest which should enhance our understanding of animal thermal ecology and improve our ability to forecast the responses of organisms to environmental change.

## Introduction

Human activities are rapidly altering the global climate, resulting in significant consequences for organisms across ecosystems (Deutsch et al., 2008; Garcia-Costoya et al., 2023; Sinervo et al., 2010; Thomas et al., 2004). In response to this challenge, considerable attention has been focused on predicting if and how organisms will adapt to changing thermal environments. To accomplish this, we must first be able to accurately quantify and map the thermal environments that organisms experience currently to forecast how these environments will change in the future. However, there are two major obstacles in accurately describing contemporary thermal environments and predicting future ones: (1) overcoming the mismatch between the spatial resolution at which environmental data are typically collected and the scale at which a particular organism experiences thermal variation, and 2) quantifying thermal landscapes without substantial measurement gaps in time or space. While these are significant challenges for all organisms, they are especially acute for ectotherms that live in the spatiotemporally heterogenous thermal environments that are typical of terrestrial habitats.

For small ectotherms that live in heterogeneous environments, interactions between an individual and its environment happen at fine temporal (minutes to hours) and spatial (sub-meter to a few square meters) scales, yet most studies to date have relied on geographically coarse environmental temperature data from sources like WorldClim or CHELSA which can only offer climatic variable estimates at scales of 30 arc-seconds (∼ 1 km^2^; Fick & Hijmans, 2017; Karger et al., 2017). Only by incorporating thermal data collected relatively continuously at the study site and at the scale of microhabitat can we capture the type of environmental variation that shapes the physiology and behavior of most terrestrial ectotherms (Bozinovic et al., 2011; Fey et al., 2019; Germain & Lutz, 2020; Sears et al., 2016; Vasseur et al., 2014; Woods et al., 2015). For example, Potter et al. (2009) demonstrated that plant leaves buffered tobacco hornworm moth (*Manduca sexta*) eggs from fatally high temperatures and variation in thermal profiles between individual leaves on a single plant caused substantial differences in development rates. Alujević et al. (2023) showed that thermoregulatory accuracy and fitness depend on variation in thermal quality at the level of individual territories (<1000 m^2^) within a single population of rock agama lizards (*Agama atra*). Many other studies have also emphasized the importance of fine-scale thermal heterogeneity in the ecology and evolution of terrestrial ectotherms (e.g., Bütikofer et al., 2020; Cox et al., 2018; Cox, Tribble, et al., 2020; Logan et al., 2014, 2015, 2016, 2021; Neel et al., 2021; Pincebourde & Suppo, 2016; Sears & Angilletta, 2015; von Schmalensee et al., 2021; Williams et al., 2022). Thus, understanding how animals interact with thermal environments in the face of climate change requires bridging the spatial gap between the broad scales at which environmental data are typically collected and the thermal ecology of the taxa of interest (Potter et al., 2013; Sears et al., 2011).

Several methods have been developed that partially overcome the challenges of measuring biologically relevant thermal environments. These include the use of biophysical (mathematical) models combined with remote sensing (where operative temperatures are predicted from biophysical theory) or physical models (data loggers) deployed at a field site (where temperatures are measured empirically; Figure S2) to assess “operative temperature distributions”, which can be thought of as null distributions of temperatures that are available to a particular organism in a given space and time. Operative temperatures are distinct from other types of temperature measurements as they represent instantaneous estimates of the equilibrium body temperature that a specific animal would attain at a given microsite (Bakken, 1992; Dzialowski, 2005). Operative temperature distributions can provide information on habitat thermal quality and the energetic requirements for the organism to perform well in that environment. Further, these distributions are usually measured at spatial scales that are much finer than those provided by publicly available climatic data. The quantification of operative temperatures has played a pivotal role in the study of animal thermal ecology, physiology, behavior, and evolution (reviewed in Angilletta, 2009), and it has been an irreplaceable tool in forecasting ecosystem responses to rapid environmental changes (Gunderson & Leal, 2012; Huey et al., 2012; Logan et al., 2013).

Although the measurement of operative temperatures has been fundamental to our understanding of animal thermal biology and predicting responses to climate change, there are limits to their use and application. For example, the biophysical modeling approach to quantifying operative temperatures requires that environmental data such as air temperature, humidity, wind speed, and solar radiation, be gathered through the deployment of weather stations or acquired from satellites, and these macroclimatic data are then integrated with corresponding spatial information, using biophysical equations to model heat exchange rates for specific organisms in distinct environments (Buckley et al., 2023; Kearney & Porter, 2017; Maclean et al., 2019; Maclean & Klinges, 2021). This approach has gained popularity in recent years thanks to the development of ecophysiological modelling tools (e.g., NicheMapR, Kearney & Porter, 2017; TrenchR, Buckely et al. 2023; microclima, Maclean et al. 2019). These tools have enabled the estimation of microclimate distributions together with predictions of body temperatures and energetic demands for a given population. While these methods have been invaluable, ecophysiological software programs face some limitations in their capacity to translate population-level estimates to the dynamics of individual organisms (Meyer et al., 2023). First, the input of high-quality environmental data is crucial, yet these data are sometimes not available at the temporal and spatial scales at which organisms typically experience their environment. Second, ecophysiological modelling usually requires general assumptions about the ways in which factors like vegetation characteristics, soil properties, and topography shape microclimates, and these assumptions can be violated in some situations (Woods et al., 2015). Third, while current software packages can model thermal conditions across a range of microhabitats, they cannot assess the presence, frequency, or spatial distribution of these microhabitats in the area of interest.

A potentially more tractable approach to quantifying operative thermal environments, particularly for small terrestrial vertebrates, involves the deployment of temperature data loggers that mimic key biophysical properties of the organism of interest (although some challenges remain; Alujević et al. 2024). These loggers are known as “operative temperature models” (OTMs) and they provide a measurement of the thermal environment at the organism’s spatial scale by integrating conductive, convective, and radiative heat transfer between the animal and its surroundings (Angilletta, 2009; Bakken, 1976; Bakken et al., 1985). While temperature distributions obtained with OTMs can be nearly temporally continuous (if OTMs are programmed to record temperatures frequently), they are spatially incomplete as OTMs can only be deployed across a small subset of locations within a given field site and thus cannot capture the full thermal complexity of spatially heterogeneous habitats. This is a problem, as the full heterogeneity and spatial structure of operative thermal environments can play a fundamental role in the thermoregulation, movement, and energetics of the organisms that live in these habitats (Sears et al., 2016; Sears & Angilletta, 2015).

Drones, or uncrewed aerial vehicles (UAVs), provide an avenue by which the challenges associated with quantifying thermal environments at biologically relevant scales might be overcome. Recent technological advancements and cost reductions of drones, in combination with their versatility and ability to access remote or challenging terrains, have led to them becoming invaluable tools in ecological research (e.g., biodiversity monitoring, Aucone et al., 2023; wildlife tracking, Saunders et al., 2022; ecosystem health assessment, Francis et al., 2022; wildfire management, Saffre et al., 2022; crop monitoring for agriculture, Cuaran & Leon, 2021; etc.). A number of commercial drone models are now outfitted with thermal imaging (thermal infrared, or “TIR”) cameras. Overlapping thermographs from transects flown by these drones, stitched together using photogrammetry, can be used to quantify the spatial structure of thermal environments across large areas at high-resolution (Thiele et al., 2017; Webster et al., 2018). Yet aerial thermography and OTMs give largely non-overlapping estimates of thermal environments. Although TIR drone-based photogrammetry can produce a thermal landscape that is comprehensive in its spatial coverage, it remains temporally discrete, producing a thermal map for a single point in time (the period during which the drone was being flown). Moreover, while OTMs integrate all the relevant forms of heat transfer present in the environment to give estimates of the equilibrium body temperatures that would be achieved by the study organism, the temperature sensors on TIR drones estimate only the heat energy that is radiatively emitted from surfaces. Factors such as the distance between the camera and the surface, the angle of incidence of the sun, ambient temperature, substrate emissivity, and wind speed (among others) can influence drone-based TIR estimates (Cilulko et al., 2013; Faye et al., 2016; Jiao et al., 2016; Kelly et al., 2019; Lathlean & Seuront, 2014; Playà-Montmany & Tattersall, 2021; Tattersall, 2016), and some TIR drone setups in some environments may require a warm-up period to reduce the impact of vignetting (Yuan & Hua, 2022). In summary, OTMs give temporally complete, but spatially discrete, distributions of operative temperatures whereas TIR drone photogrammetry gives spatially complete, but temporally discrete distributions of emitted surface temperatures. To achieve the goal of measuring and predicting spatiotemporally complete distributions of operative temperatures, one must correct the drone-based temperature estimates such that they describe operative temperatures and then extrapolate those temperatures to days and times when photogrammetry data are not available.

Here, we present a new method that integrates TIR drone photogrammetry with field-deployed OTMs to generate fine-scale, spatiotemporally complete landscapes (maps) of operative temperature for terrestrial organisms. At its core, our method takes biologically relevant and temporally continuous, but spatially limited, OTM data and extrapolates them to a broad geographic area at high-resolution. We developed a new R package called `throne (`‘`th`’ermal d‘`rone`’)that streamlines our method and renders it accessible to biologists and conservation managers by requiring minimum user input to produce high-resolution operative temperature maps for any day/time combination during which OTMs were deployed. Finally, we used operative temperature data for the western fence lizard (*Sceloporus occidentalis*) to validate our method at a thermally complex field site in the Great Basin Desert of northern Nevada. Our method for quantifying and predicting terrestrial thermal environments at extremely high spatiotemporal resolution has the potential to increase our understanding of how individuals, and thus populations, will respond to climate warming, habitat conversion, and other environmental changes.

## throne workflow

Our approach takes biologically relevant and temporally continuous, but spatially limited, OTM data and extrapolates them to a broad geographic area at high-resolution by leveraging the power of TIR drone photogrammetry. The general workflow involves the following steps: 1) Simultaneously collecting operative and emitted surface temperatures (hereafter, “surface temperatures”) using OTMs and a TIR compatible drone, respectively; 2) Integrating these datasets and applying necessary corrections; and 3) Interpolating corrected data across time and space to generate “predicted” thermal landscapes (Figure 1). The user first assembles a map of surface temperatures of their field site using TIR drone photogrammetry while at the same time collecting operative temperatures using OTMs deployed at microsites that capture local microhabitat diversity. All subsequent steps can be accomplished using our R-package, ‘`throne`’, available on GitHub (https://github.com/ggcostoya/throne). These datasets are then combined such that the relationship between OTM temperatures and drone surface temperatures is assessed and necessary corrections are applied. Finally, OTMs are assigned to tiles (i.e., unique latitude and longitude combinations) from the thermal orthomosaic with similar thermodynamics and a high-resolution operative temperature map is then generated for any day and time during which the OTMs were logging. The assembly of thermal landscapes is automated and only requires operative and surface temperature data (with corresponding metadata) and a small number of parameters (e.g., level of smoothing of temporal thermal profiles and desired spatial resolution, date and time for the final thermal landscape; Figure 2) as input. We detail each of the steps below.

**Figure 1.**
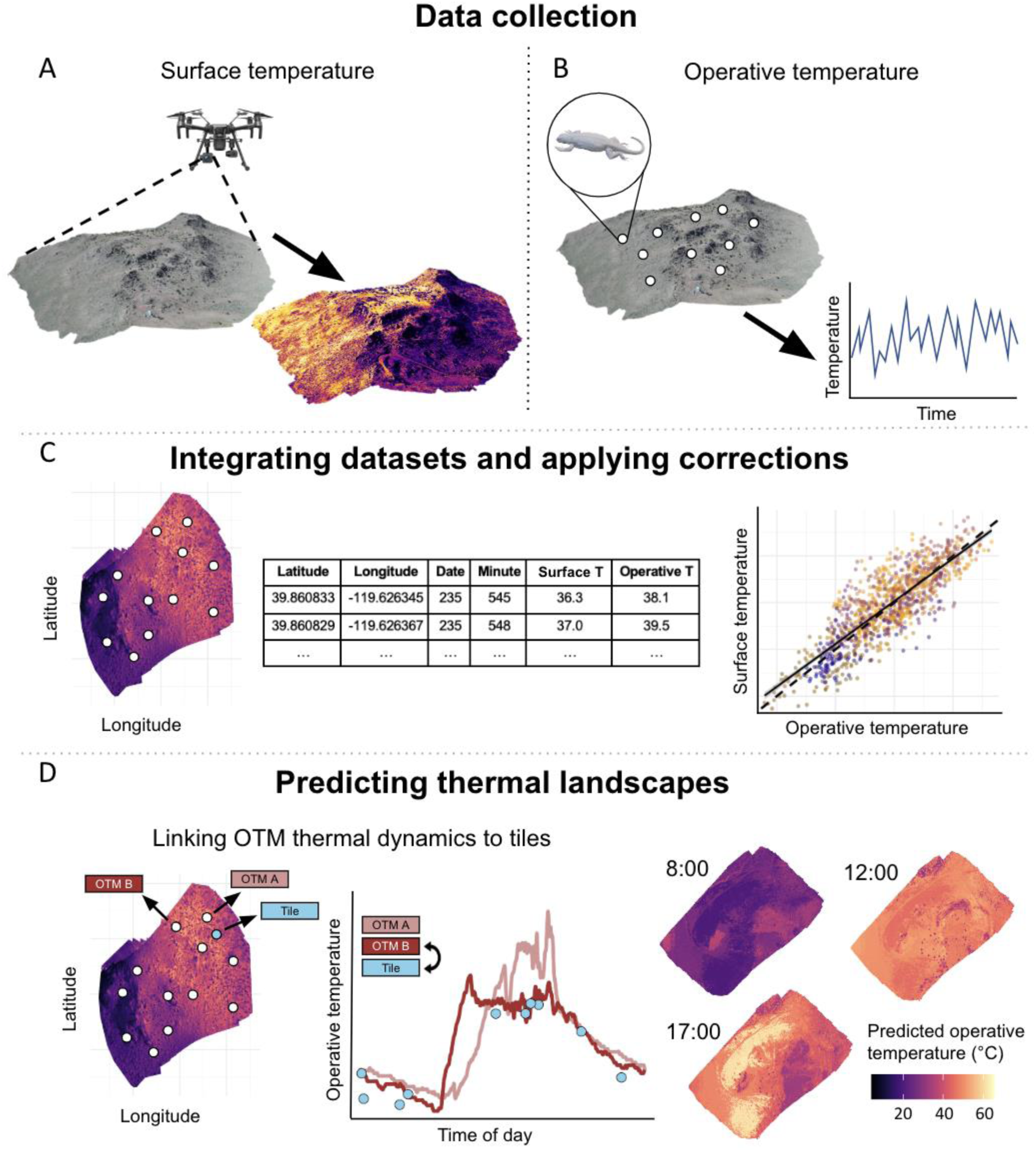
General workflow for our method that links TIR drone photogrammetry with operative temperature data to produce spatiotemporally complete and biologically relevant thermal landscapes. The practitioner or researcher first assembles a map of surface temperatures of their field site using TIR drone photogrammetry (A) while at the same time collecting operative temperatures using OTMs matching the biophysical properties of their study organism and that are deployed across a range of microsites that capture a representative sample of local microhabitat diversity (B). All subsequent steps can be accomplished in our R package, `throne`. C) These datasets are then combined such that the relationship between OTM temperature and drone-based surface temperature is assessed and necessary corrections are applied. D) The thermal dynamics of the OTMs are assigned to tiles from the thermal orthomosaic and a high-resolution operative temperature map can then be generated for any day and time during which the OTMs were logging.

**Figure 2.**
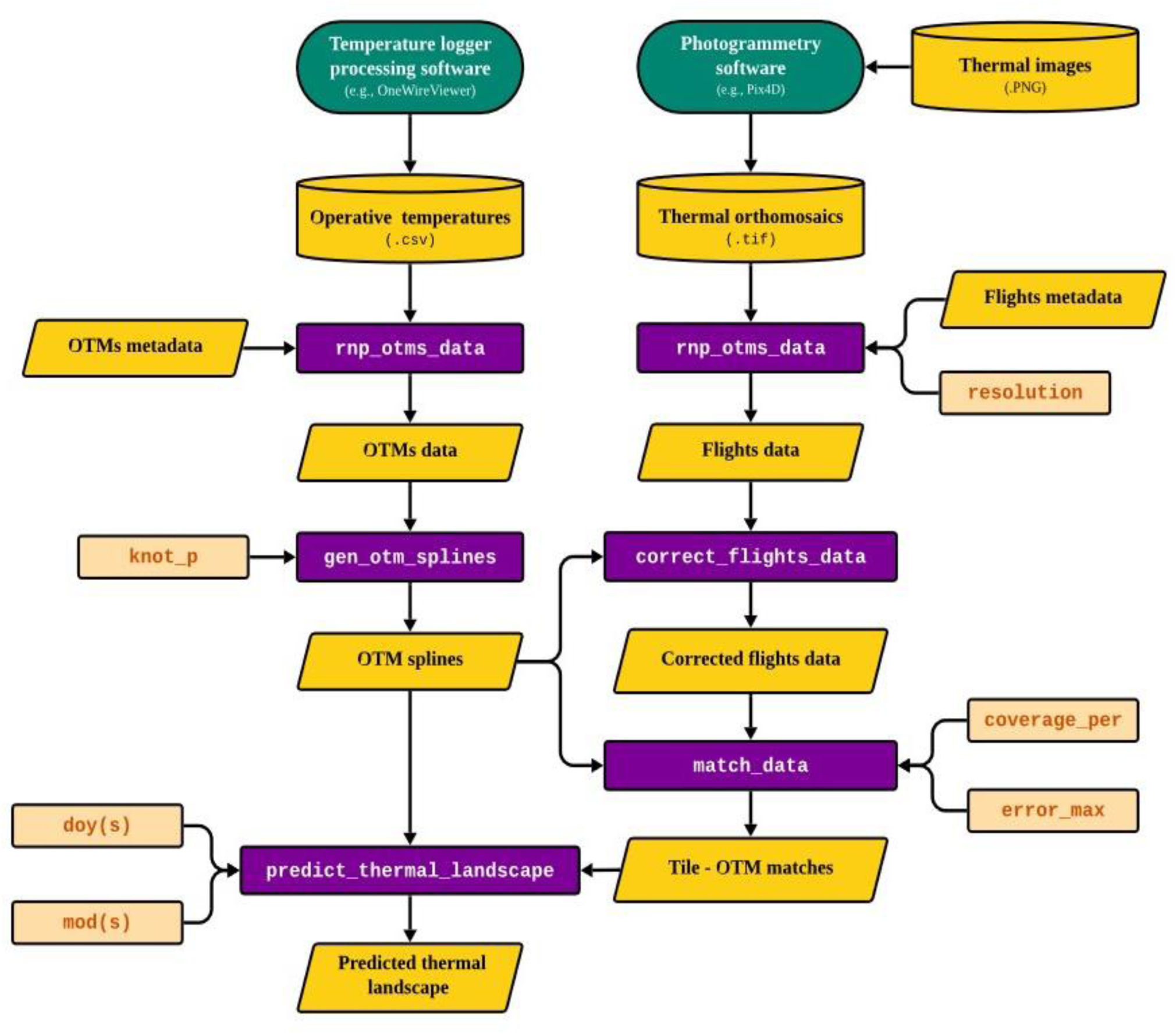
The `throne` R-package workflow. Sources of input data are represented by yellow cylinders for databases and yellow tilted squares for single datasets. Green ovals denote software tools external to `throne.` Squares with red font represent user-specified parameter values. Purple rectangles denote embedded functions.

### 1. Data collection

#### 1.1. OTM deployment

OTMs should be built to mimic a key set of biophysical properties of the organism of interest (e.g., shape, surface reflectance, conductivity), ensuring that under constant environmental conditions they produce an “instantaneous” estimate of the equilibrium body temperature that the animal would achieve at the microsites in which the OTMs are deployed. OTMs can be built using different techniques that have varying degrees of practicality and accuracy, although recent advancements in 3D printing can be used to generate highly accurate and cost-effective OTMs for many species (Alujević et al., 2024). For our method to work optimally, OTMs should be deployed strategically across different microhabitats and physiographic features (e.g., substrate types, exposure level, vegetation types, slopes) within the study area to capture a wide range of microclimatic conditions.

#### 1.2. Drone flights

Thermal photogrammetry data are collected using a drone equipped with an IR thermal imaging camera. A flight plan can be programmed into the drone using photogrammetry software, and this software sometimes comes with the drone when purchased. For our method to be most successful, the following guidelines should be followed: 1) Flights should be conducted under weather conditions that maximize thermal heterogeneity (e.g. sunny weather) as this increases the ability of `throne` to match OTMs with tiles of similar thermodynamics; 2) Flights should be distributed across different days and times to maximize representation of daily temperature fluctuations, although nighttime flights are likely not necessary because thermal heterogeneity is typically much lower without solar radiation such that flights near dawn and dusk will be sufficient to capture these dynamics; 3) To prevent stitching issues, flights should be conducted over a slightly larger area than the specific area of interest; 4) Flight times should be kept relatively short to ensure that all images are captured during similar site-level thermal conditions and within an ecologically relevant window; 5) The resolution of the mounted thermal imaging camera should be considered when planning flights and choosing flight altitude; 6) The mounted thermal imaging camera should be radiometrically calibrated and produce thermal images in either TIF or R-JPEG file formats; 7) The average emissivity of the site substrate should be assessed to appropriately calibrate the thermal imaging camera; 8) The vertical and horizontal percentage of overlap between photos should be set to a high value (preferably 90% or greater but no less than 70%). Image overlap reduces the measurement error inherent in individual IR images; 9) Ground control points (GCPs) should be deployed as they help to accurately georeference thermal images and minimize processing errors (GCPs should be strategically placed to cover the study site’s borders and topographical complexity). After collecting thermal images, photogrammetry software can be used to process images and generate a thermal orthomosaic—a composite snapshot of the thermal landscape. Image processing steps will vary by software type (for more details on this process see the `throne` website: https://ggcostoya.github.io/throne/), although the resulting raster file should be produced in the TIF-file format.

### 2. Data processing

#### 2.1. Operative temperatures from OTMs

After the OTMs are retrieved from the field and the individual OTM data are downloaded, the OTM data files are then combined into an R data frame, using the `rnp_otms_data` function in `throne`. This function returns a processed data frame with columns for the OTM’s identifier, year, day of the year, minute of the day, and operative temperature where each row is a unique operative temperature measurement at a given time by a given OTM. Further, rnp_otms_data incorporates user-specified metadata in the final output, including the geographic position where the OTM was deployed which, if needed, is projected into Universal Transverse Mercator (UTM) coordinates to ensure compatibility with flights data. Next, the output of the `rnp_otms_data` function is processed using the `gen_otm_spline` function to fit smoothing spline models to each individual OTM for each day during its deployment in the field using the base R native `smooth.spline` function (R Core Team 2024). Briefly, these models generate a smoothing function that captures the essential thermodynamics of each OTM throughout a given day. Smoothing minimizes noise from short-term, stochastic shifts in operative temperature while capturing fluctuations caused by conditions unique to each day (e.g., sustained changes in cloud cover or wind) that may be ecologically relevant. This approach eliminates the need to collect additional metadata during OTM deployment (e.g., aspect or shade cover) and removes the assumption that within-day thermal fluctuations follow a fixed sinusoidal curve in order to inform the resulting models. Nonetheless, as the level of smoothing may be important in certain systems and may influence the ability of `throne` to match OTMs with orthomosaic tiles, the `gen_otm_splines` function incorporates the parameter `knot_p`. This parameter determines the percentage of observations recorded by an OTM in a given day that are used to determine the number of knots in the spline model (fewer knots equate to more smoothing). The output of the `gen_otm_splines` function inherits all metadata information from the OTM data frame while adding a nested column containing the spline model. The resulting data frame contains as many rows as there were combinations of unique OTM identifier and date as each of these combinations is assigned a fitted spline model.

#### 2.2. Surface temperatures from drone flights

Drone flight orthomosaic `.tif` files generated using photogrammetry software are converted into an R data frame using the function `rnp_flights_data`. File .tif format is a standard for thermal raster images, containing pixel-wise temperature values generated from radiometrically calibrated sensors (note that file formats like JPEG are not suitable inputs for our approach as they do not preserve the absolute temperature data needed for accurate thermal predictions). The rnp_flights_data function first reads the `.tif` file as a `raster` within the R environment using functionality from the package terra (Hijmans et al., 2024). Second, the function summarizes the data to the desired spatial resolution via the argument `resolution` . The resolution argument determines the area covered (in m^2^) by each of the orthomosaic tiles of the final output and it can be set to any value ≥ 0.5 m^2^. Lastly, the function adds user-specified metadata to the processed flight dataset including values for year, day-of-year, and minute of the day at which the flight started and ended. The final output is a data frame with columns for geographic position indicated by an “x” and “y” coordinate within the UTM zone where the flight took place, time (year, day-of-year, minute of the day at which the flight started and ended) and surface temperature. In this data frame, each row corresponds to the surface temperature in a given tile (unique x-y combination) of an individual flight.

### 3. Integrating OTM and drone data

#### 3.1. Converting surface temperatures to operative temperatures

Due to the fundamental differences in the physical properties of surface (IR-based) and operative (OTM-based) temperature measurements, the data frames representing thermal maps obtained via the `rnp_flights_data` function need to be corrected such that they represent operative temperatures. To achieve this, `throne` includes the `correct_flights_data` function. First, the function identifies all tiles within the study area that contained OTMs, and 1) gathers all the temperature measurements of those tiles collected across multiple flights, and 2) estimates the temperature experienced by all OTMs at the exact set of dates and times when each of the flights took place using the spline models obtained via the `gen_otm_splines` function. Second, the function estimates the average bias between surface and operative temperature measurements for each flight (Figures S5 and S6) and subtracts this bias from all surface temperature measurements. This first correction step is needed because the magnitude of the difference between the OTM and surface temperatures can vary systematically between days and by time-of-day due to the difference in the way these different measurements respond to ambient temperature, solar angle, overall light availability, etc. However, the function offers users the ability to choose which metric they prefer as the basis for surface temperature correction (average, median, or mode), or the option to skip the correction step entirely. Lastly, the function inspects the relationship between surface temperature (which the user may or may not have chosen to correct for date and time-of-day) and operative temperature by fitting a simple linear regression between these variables. If temperature measurements collected by the drone were perfectly unbiased estimates of operative temperature, the relationship between these two variables would be 1:1 (intercept = 0 and slope = 1). To achieve this 1:1 relationship, the function applies a second correction following the equation:

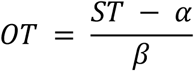

Where ST and OT are surface and operative temperatures respectively, *α* and *β* are the intercept and slope of the relationship between ST and OT across all tiles where OTMs were deployed. To give further control over how this correction step is performed, correct_flights_data also offers users the possibility of applying this step while accounting for day- and time-specific relationships. In this case, the correction is applied via the equation:

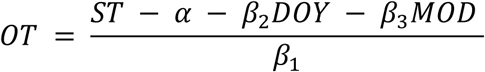

Where *β*_1_ is the effect of OT on ST (same parameter as *β* in the previous equation) while *β*_2_ and *β*_3_ are the effects of day of the year (DOY) and median minute of the day when the flight took place (MOD) respectively.

### 4. Predicting thermal landscapes

The last step of the `throne` workflow is generating a thermal landscape for a specific time of interest. This is achieved via the `match_data` and `predict_thermal_landscape` functions. First, the `match_data` function matches each tile in the thermal orthomosaic with an OTM based on the similarity in their thermodynamics. For each tile, the function calculates the average absolute difference between the temperatures recorded in that tile across multiple flights (previously corrected through the `correct_flights_data` function), and the temperatures extracted from all OTM splines at the same time the flight took place. OTMs do not need to have logged temperatures at the exact time the flight took place as the operative temperatures are predicted from the date specific OTM spline function (`gen_otm_splines`). To assign a match to a tile, the `match_data` function chooses the OTM that minimizes the absolute difference between temperature measurements. For some tiles, it is possible that their thermodynamics are still notably different than those of the OTM that best describes it among the OTMs deployed (i.e., that the average absolute error is substantially large). To account for that, users can specify the maximum error they are willing to tolerate for a tile’s thermal profile to be included in their final output by setting the `error_max` parameter (if the minimum error is larger than `error_max`, the pixel is not assigned an OTM). If it is not specified, error_max will default to 100, a sufficiently large number to permit any match, which in turn allows users to explore the quality of their matching (Figure S13) and perform any diagnostics or quality control efforts they see fit. The resulting output of the `match_data` function is a data frame where each tile is associated with a character indicating the identifier of the OTM that best describes its thermodynamics. Finally, the `predict_thermal_landscape` function takes the output of the data frame obtained through `match_data` and calculates the predicted operative temperature in each tile at a user-specified date and time using the OTM and date specific cubic spline models fitted through the `gen_otm_splines` function. Through this approach, users can predict a thermal landscape for any date and time (minute of the day) within the general period when OTMs were logging (i.e., for all days in which there are fitted spline models). The final output is a data frame with six columns, indicating the year, day of year, minute of the day, x and y UTM coordinates, and predicted operative temperature for that specific tile.

#### Validation

To validate our approach to mapping operative thermal environments, we tested 1) whether the predicted thermal landscapes output by `throne` are accurate for microsites where OTMs were not deployed, and 2) the sensitivity of the method to different user choices (e.g., number of drone flights, number of OTMs, level of spline smoothing). To accomplish this, we conducted two validations where we compared the operative temperatures predicted by `throne` with those recorded by OTMs that had been withheld from the workflow. With our validations we demonstrate how changing the number of flights conducted, OTMs deployed, and spatial and temporal scale affect the accuracy and utility of our method. Validation 1 was conducted over a 76-day period (May 15th to July 29th, 2023) and for a ∼33,000 m² study area, with 10 flights and 73 OTMs. In contrast, validation 2 took place over 3 days within a smaller area (600 m²) but involved a higher frequency of flights (34). For each validation, we executed the complete `throne` pipeline with different combinations of parameters (subset of flights, subset of OTMs, and `knot_p` values). Below, we detail how we collected data to conduct each validation in the context of our study system and provide more details on the methodology we followed to conduct these validations.

## Methods

### 1. Data collection

#### 1.1. Validation 1

##### 1.1.1. OTM deployment

We deployed 73 operative temperature models (OTMs) on April 15th, 2023, across a 33,000 m^2^ area southwest of Pyramid Lake in Washoe County, Nevada, USA (39.865 N, 119.624 W). This site is in the Great Basin Desert and is characterized by a thermally complex array of ridges and rocky outcrops (Figure S1). The vegetation community is dominated by big sagebrush (*Artemisia tridentata*), saltbush (*Atriplex gardneri*), pinyon pine (*Pinus monophylla*) and juniper (*Juniperus osteosperma*). We 3D printed our OTMs using acrylonitrile butadiene styrene (ABS) for studies of the thermal ecology of the western fence lizard (*Sceloporus occidentalis*; Figure S2), a species that is abundant at our field site, and for which we previously validated 3D printed OTMs against live lizards in the field (Alujević et al., 2024). We centrally suspended a temperature logger (iButton, accuracy of 0.5 °C, Embedded Data Systems, Lawrenceburg, KY) inside each model and set each logger to record temperature every 50 minutes (∼ 29 measurements / day). We obtained operative temperature measurements from the loggers for 119 days from April 16th to August 19th excluding June 11th, 12th and 27-29^th^ (these were days when we had to retrieve OTMs to download data due to the limited storage capacity of iButtons). This time period and geographic area encompassed significant seasonal and spatial thermal variation (Figure S3).We distributed OTMs across microhabitats that are typical of this site (for detailed descriptions of these microhabitats, see Table S1) and at different orientations to capture a range of ecologically relevant microsites based on our previous experience working with this species in this environment. For each OTM, we recorded its position in a World Geodetic System projection (WGS; i.e., latitude and longitude) using a high accuracy (< 1 m^2^ resolution) Trimble Geo7x handheld GPS unit (Trimble, Westminster, CO).

##### 1.1.2. Drone flights

We created a flight mission using DJI Pilot software (v 1.1.5; DJI, Shenzhen, China) that was larger (52,000 m^2^) than the area covered by our OTMs to ensure that the central area of interest was fully covered by the drone transects. We placed four ground control points (GCPs) within the borders of the OTM deployment area, covering approximately the highest, middle, and lowest elevation (Agüera-Vega et al., 2017). We recorded each GCP’s position in WGS projection using the same Trimble Geo7x handheld GPS unit. The flight altitude was set at 100 m above ground level (AGL), resulting in a ground sample distance (GSD) of 13.08 cm/pixel. The mission flight speed was 5.3 m/s, with a 70% side to 80% frontal image overlap ratio. Thermal and RGB images were collected simultaneously in-flight using a FLIR Zenmuse XT2 infrared camera (focal length = 13 mm, TIR resolution = 640 × 512, spectral range = 7.5-13.5 μm, accuracy = <50 mK @ f/1.0, file format = TIFF/R-JPEG/JPEG, and emissivity = 1.0) mounted on a DJI Matrice 200 Series V2 quadcopter. We conducted 10 mission flights between 8:30 AM and 7:45 PM in the period between May 15th to July 29^th^, 2023 (Table S2). The average wind speed (±SD) across all flights was 3.9 ± 4.3 m/s and the average air temperature was 23.5 ± 11.5 °C. We flew 8 of these missions under sunny and/or clear conditions whereas two occurred during light cloud cover, conditions that are representative of the typical weather experienced at our field site during summer months.

We processed all drone imagery using the Thermal Camera processing template in Pix4Dmapper (version 4.8.4; Pix4D; Prilly, Switzerland). We included both thermal and RGB images in the template with the following changes to the default settings for thermal image processing: The Point Cloud Point Density was set to ‘high’ and the orthomosaic was generated as a GeoTIFF with ‘merge tiles’ enabled. We georeferenced all images using GCPs to ensure maximum accuracy. We successfully generated 34 orthomosaic raster images and created a `.tif` file for each of the processed flights, with an RMS error (the difference between the initial and computed positions of the GCPs) of 0.0475 ± 0.0063 m (mean ± SD) and an average orthomosaic GSD of 4.44 ± 0.474 cm/pixel. To test for the effect that using GCPs had on the accuracy of the final thermal landscape output of `throne`, we generated an alternative set of orthomosaic raster images during which we skipped the georeferencing step.

#### 1.2. Validation 2

##### 1.2.1. OTM deployment

We deployed 33 OTMs from August 24 -26, 2023, across a 600 m^2^ area in Washoe County, Nevada, USA (39.868 N, 119.627 W; the same general field site as in validation 1), logging temperature data every 2 minutes, following the methods described above.

##### 1.2.2. Drone flights

We created a flight mission over a 1500 m^2^ area that encompassed the OTMs and three GCPs that were strategically deployed within the area as described in validation 1. The flight altitude was set at 40 m AGL, resulting in a GSD of 5.23 cm/pixel. We conducted 34 mission flights between 8 AM on August 24 to 7 PM on August 26, 2023 (Table S2). The mission flight speed was 2 m/s, with a 70% side to 80% frontal image overlap ratio. The average wind speed across all flights was 3.8 ± 3.1 m/s and the average air temperature was 27.0 ± 5.1 °C. We flew 30 of these missions under sunny and/or clear conditions whereas four occurred during light cloud cover, conditions that are representative of the typical weather experienced at our field site during summer months. We successfully generated 34 orthomosaic raster images and created a .tif file for each of the processed flights, with an RMS error of 0.0807 ± 0.0035 m (mean ± SD) and an average orthomosaic GSD of 2.35 ± 0.69 cm/pixel.

### 2. Data processing

After the iButtons were retrieved from the field for each validation, we downloaded the data using OneWireViewer software (Analog Devices, Inc., Wilmington, MA) and ran the data through the `throne` workflow. We used the `rnp_otms_data` function to process all raw OTM data into a single data frame which we later used to fit OTM and date-specific cubic splines using the `gen_otm_splines` functions. To test how the choice of smoothing parameter (`knot_p)` affected the accuracy of our predictions, we fitted cubic splines with three different knot_p values for each validation (see below). Flight-specific orthomosaic `.tif` files (both GCP-referenced and not) were imported into R and converted into data frames using the function `rnp_flights_data`. For the latter function, we set the `resolution` to 1, resulting in a spatial resolution of 1 m^2^ for the thermal landscape, an area that is ecologically relevant to western fence lizards based on their body size and home range as determined by our research group previously (unpublished data) and by studies in other populations in the western USA (Davis & Ford, 1983; Sheldahl & Martins EP, 2000).

### 3. Predicting thermal landscapes

To validate that the final thermal landscapes produced by our method can accurately estimate operative temperature in areas of field sites where OTMs were not deployed (Garcia-Costoya, unpublished data), we generated a predicted thermal landscape from a subset of our deployed OTMs and then compared predicted operative temperatures to actual (observed) operative temperatures for the tiles that contained the remaining OTMs. We did this separately for each validation and examined how three parameters influenced `throne`’s predictive accuracy: number of drone flights, number of OTMs deployed, and the `knot_p` smoothing parameter. We assessed the number of flights and OTMs required to produce accurate thermal landscapes as these factors are likely the most important from a logistical and budgetary standpoint and therefore may reduce the usefulness of our method for some practitioners. We were also interested in examining the effect of `knot_p` as this parameter will determine the sensitivity of our method to short term fluctuations and thus our ability to accurately predict highly temporally variable landscapes. To examine how variation in these parameters affects the accuracy of the final predicted thermal landscape we ran the entire `throne` workflow a total of 52 times per validation, predicting operative temperature landscapes using different numbers of drone flights (validation 1: 3, 5, or 10; validation 2: 3, 8, 17 or 34), OTMs deployed (validation 1: 10, 30 or 70; validation 2: 10, 20 or 33), and knots for the spline models (validation 1: 0.25, 0.5 or 1; validation 2: 0.017, 0.067, 0.133, 0.5) which are equivalent to 0.3, 0.6, and 1.2 knots/h, respectively, for validation 1 and 0.5, 2, 4, and 15 knots/h, respectively, for validation 2). The individual drone flights and OTMs used to evaluate each combination of parameters were randomly sampled and we replicated the test ten and five times per combination of parameters for validation 1 and 2, respectively. In the case of the drone flights, they were separated between being conducted in the morning (before 10:00), middle of the day (between 10:00 and 16:00) and evening (after 16:00) and sampled evenly across these three categories. For each combination of parameters, we calculated the differences between the actual OTM measurements and the predicted operative temperatures from the thermal landscapes for 100 random combinations of dates and times for the period during which OTMs were deployed.

## Results

### 1. Integration of OTM and drone data

Both of our validations, whether done over a longer period of time with lower frequency of flights or over a shorter period of time with higher frequency of flights, demonstrated that our method produces similar levels of high accuracy irrespective of the spatial or temporal scale of environmental sampling (Table 1).

**Table 1.**
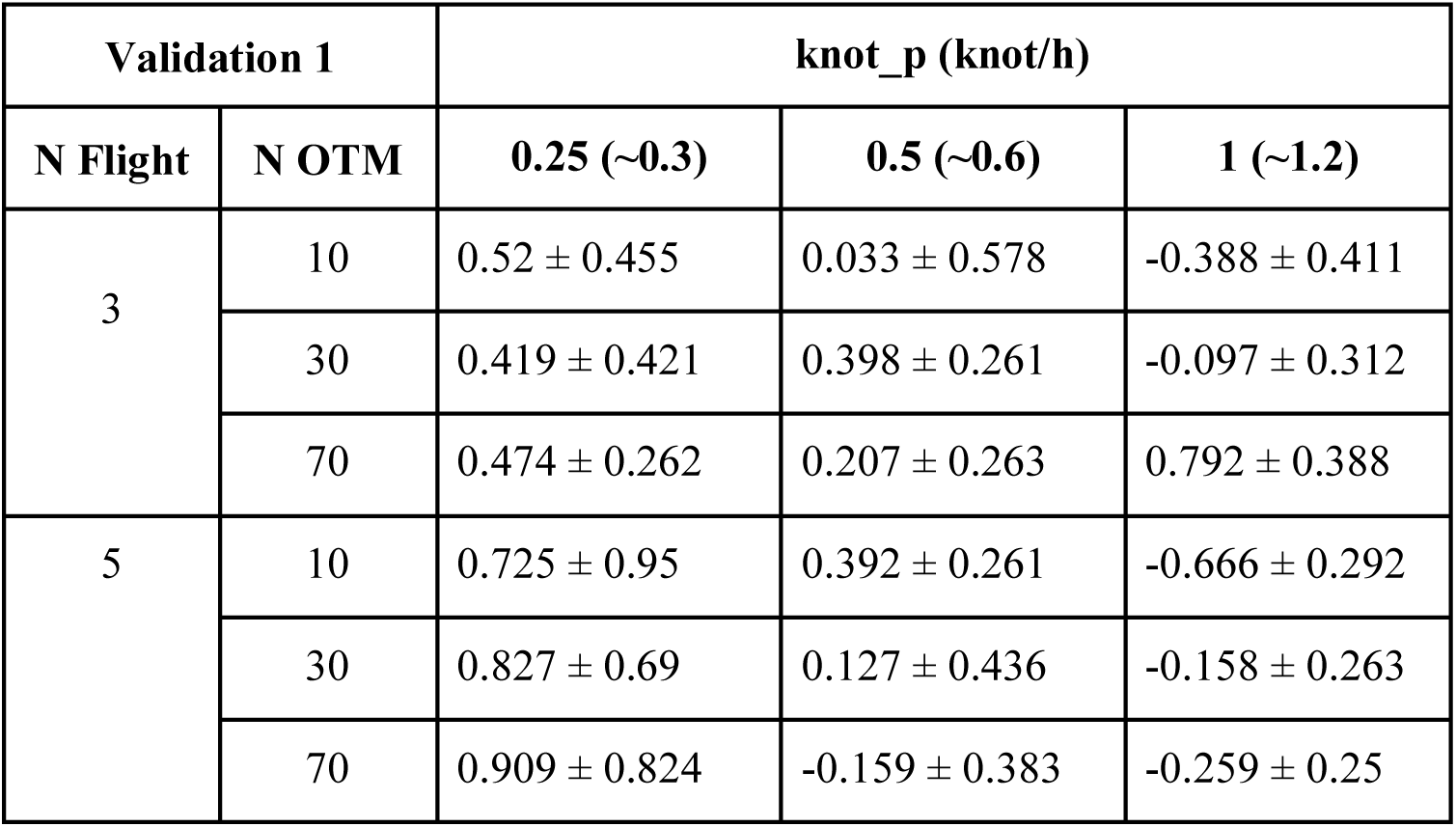
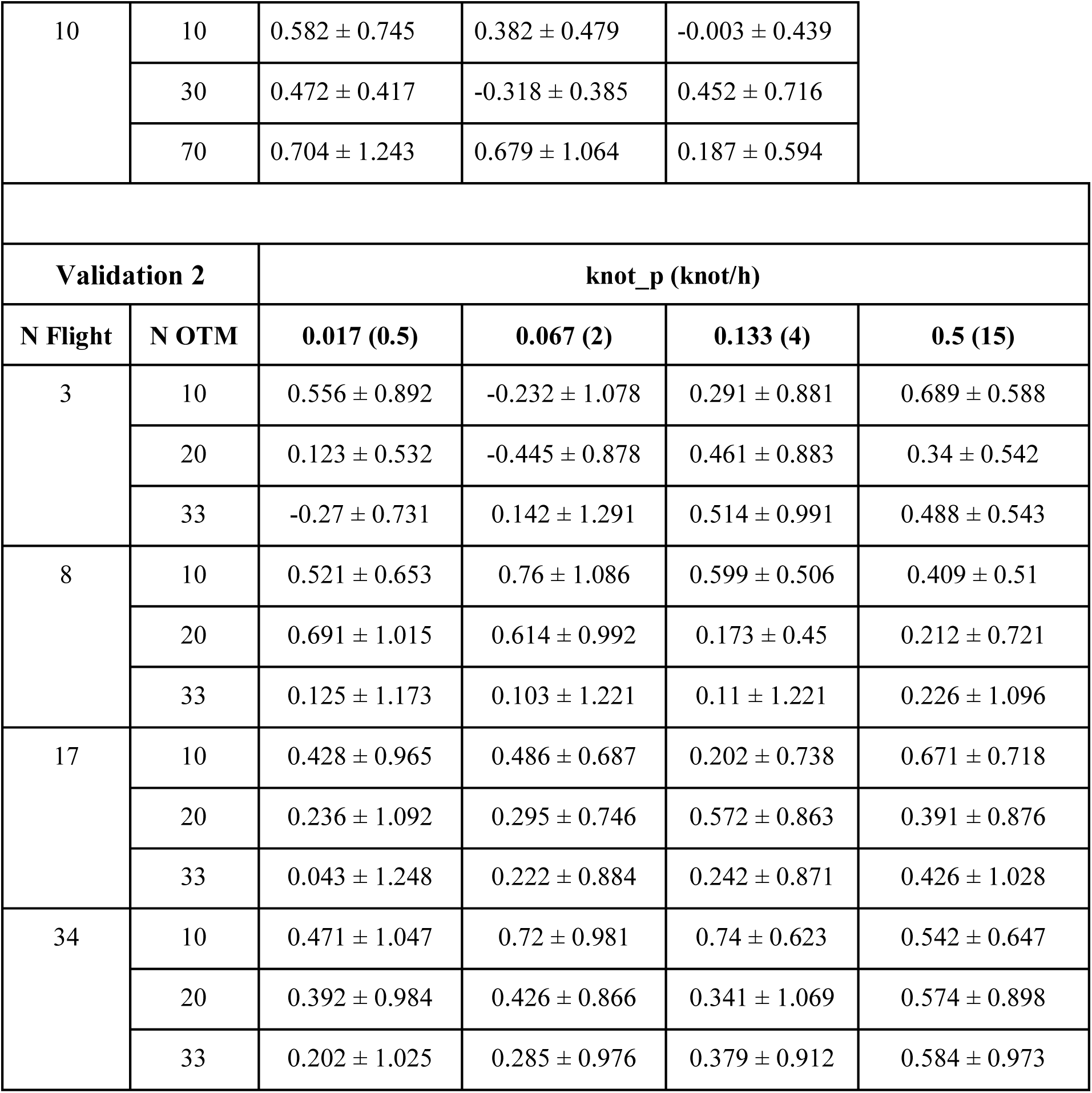
Mean predictive error (± SD) for validation 1 and 2, presented as the relative difference between the observed (from OTMs) and predicted (from the final thermal landscape output by throne) temperatures across different combinations of number of drone flights, number of OTMs, and knot_p values (magnitude of smoothing of raw OTM data; knots per hours are given in the brackets) used to generate the final thermal landscape. All comparisons were done for data collected during daytime hours (7 AM to 7 PM).

| Validation 1 |  | knot_p (knot/h) |  |  |
| --- | --- | --- | --- | --- |
| N Flight | N OTM | 0.25 (~0.3) | 0.5 (~0.6) | 1 (~1.2) |
| 3 | 10 | $0.52 \pm 0.455$ | $0.033 \pm 0.578$ | $-0.388 \pm 0.411$ |
| | 30 | $0.419 \pm 0.421$ | $0.398 \pm 0.261$ | $-0.097 \pm 0.312$ |
| | 70 | $0.474 \pm 0.262$ | $0.207 \pm 0.263$ | $0.792 \pm 0.388$ |
| 5 | 10 | $0.725 \pm 0.95$ | $0.392 \pm 0.261$ | $-0.666 \pm 0.292$ |
| | 30 | $0.827 \pm 0.69$ | $0.127 \pm 0.436$ | $-0.158 \pm 0.263$ |
| | 70 | $0.909 \pm 0.824$ | $-0.159 \pm 0.383$ | $-0.259 \pm 0.25$ |

|  |  |  |  |  |  |
| --- | --- | --- | --- | --- | --- |
| 10 | 10 | $0.582 \pm 0.745$ | $0.382 \pm 0.479$ | $-0.003 \pm 0.439$ | |
| | 30 | $0.472 \pm 0.417$ | $-0.318 \pm 0.385$ | $0.452 \pm 0.716$ | |
| | 70 | $0.704 \pm 1.243$ | $0.679 \pm 1.064$ | $0.187 \pm 0.594$ | |
| **Validation 2** |  | **knot_p (knot/h)** |  |  |  |
| **N Flight** | **N OTM** | **0.017 (0.5)** | **0.067 (2)** | **0.133 (4)** | **0.5 (15)** |
| 3 | 10 | $0.556 \pm 0.892$ | $-0.232 \pm 1.078$ | $0.291 \pm 0.881$ | $0.689 \pm 0.588$ |
| | 20 | $0.123 \pm 0.532$ | $-0.445 \pm 0.878$ | $0.461 \pm 0.883$ | $0.34 \pm 0.542$ |
| | 33 | $-0.27 \pm 0.731$ | $0.142 \pm 1.291$ | $0.514 \pm 0.991$ | $0.488 \pm 0.543$ |
| 8 | 10 | $0.521 \pm 0.653$ | $0.76 \pm 1.086$ | $0.599 \pm 0.506$ | $0.409 \pm 0.51$ |
| | 20 | $0.691 \pm 1.015$ | $0.614 \pm 0.992$ | $0.173 \pm 0.45$ | $0.212 \pm 0.721$ |
| | 33 | $0.125 \pm 1.173$ | $0.103 \pm 1.221$ | $0.11 \pm 1.221$ | $0.226 \pm 1.096$ |
| 17 | 10 | $0.428 \pm 0.965$ | $0.486 \pm 0.687$ | $0.202 \pm 0.738$ | $0.671 \pm 0.718$ |
| | 20 | $0.236 \pm 1.092$ | $0.295 \pm 0.746$ | $0.572 \pm 0.863$ | $0.391 \pm 0.876$ |
| | 33 | $0.043 \pm 1.248$ | $0.222 \pm 0.884$ | $0.242 \pm 0.871$ | $0.426 \pm 1.028$ |
| 34 | 10 | $0.471 \pm 1.047$ | $0.72 \pm 0.981$ | $0.74 \pm 0.623$ | $0.542 \pm 0.647$ |
| | 20 | $0.392 \pm 0.984$ | $0.426 \pm 0.866$ | $0.341 \pm 1.069$ | $0.574 \pm 0.898$ |
| | 33 | $0.202 \pm 1.025$ | $0.285 \pm 0.976$ | $0.379 \pm 0.912$ | $0.584 \pm 0.973$ |

Raw surface temperatures obtained from the drone (whether or not images were georeferenced using GCPs) were positively and significantly correlated with operative temperatures obtained from OTMs (p < 0.001 for both cases). However, the relationship between surface and operative temperatures obtained from non-georeferenced images was noticeably worse (intercept = -0.28, slope = 0.9, R^2^ = 0.562, df = 341) than that obtained from georeferenced images (intercept =-0.14, slope = 0.87, R^2^ = 0.62, df = 351; Figure S4). The correction process implemented via the `correct_flights_data` function (including correcting for day and time-of-day bias; Figure S5) resulted in further improvement of the fit between OTM and surface temperatures while minimally impacting the form of the relationship (intercept = 0, slope = 1, R^2^ = 0.61, df = 351; Figure S4).

When using all flights and OTMs, and a `knot_p` value of 0.5 (equivalent to 0.6 knot/h at a sampling rate of 1.2 OTM measurements per hour) to predict the full thermal landscape, half of the thermal variability across our field site was explained by as few as five OTMs, with 24 OTMs explaining 90% of the overall thermal variability (Figure 3). OTMs that were deployed at similar orientations (e.g., those deployed on surfaces that faced a particular cardinal direction) tended to be matched with spatially aggregated clusters of tiles which were in the parts of the habitat that faced those directions (Figure 3). We show predicted thermal landscapes generated for our study site in Figure S7.

**Figure 3.**
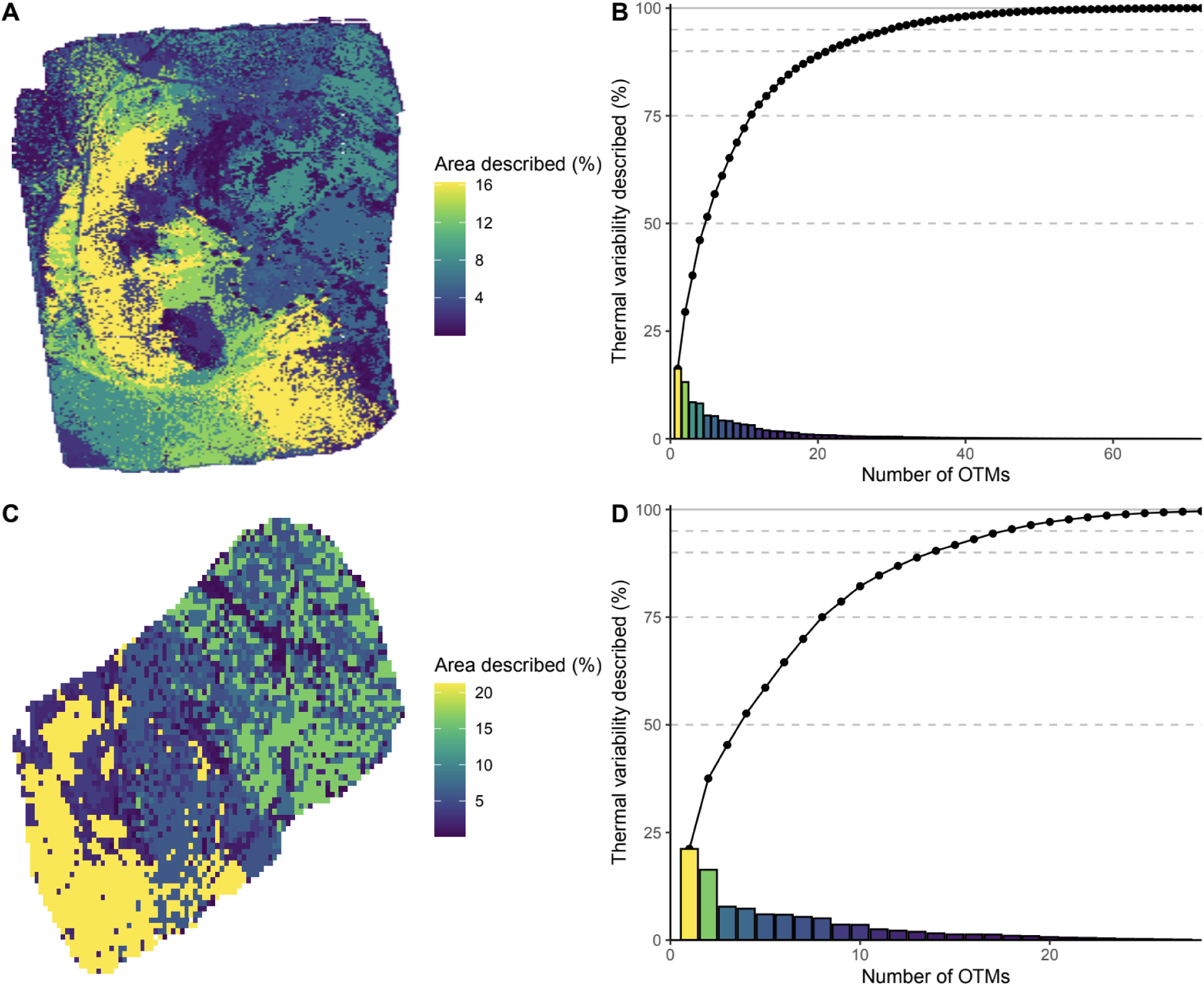
A small number of OTMs described most of the thermal variation within our field site, whether we used our method to predict operative temperatures over a longer span of time (76 days) and a larger geographic area (33,000 m^2^; validation 1; A and B), or over a shorter span of time (3 days) and smaller geographic area (600 m^2^; validation 2; C and D). The solid lines in the panels on the right show the cumulative thermal variation explained by a given number of OTMs. 22 OTMs described more than 90% of the thermal variation for the larger area (B), whereas 15 OTMs described the same amount of thermal variation for the smaller area (D).

### 2. Predictive accuracy

When we used 70 OTMs, 10 flights, and set `knot_p` to 1 (i.e., 1 knot / hour) to generate the predicted thermal landscape, the mean predictive error (the difference between predicted and observed operative temperature) was 0.187 ± 0.594°C (mean ± SD). When only 10 OTMs and 3 flights were used to generate the predicted thermal landscape (also with `knot_p` set to 1), the mean predictive error was -0.388 ± 0.411°C. In all cases, the 95% confidence interval around the mean predictive error overlapped with zero. A full list of mean predictive errors (± SD) for all combinations of flight number, OTM number, and number of knots per hour are presented in Table 1, S3, and S4 (reported as both relative and absolute predictive error). Frequency distributions of predictive errors across different combinations of parameters are shown in Figures S8-12.

## Discussion

The traditional approaches to measuring terrestrial thermal environments and predicting how they will change in the future face challenges due to limited availability of high-resolution environmental data. Our method, which integrates drone infrared imaging with *in-situ* OTM measurements, is a tractable approach to measuring terrestrial thermal environments at high spatiotemporal resolution. Importantly, our method generates thermal maps that are biologically relevant to the study species, at least to the extent that OTMs are properly designed and calibrated for the organism of interest. We have made our method accessible to researchers and practitioners by developing an R package, `throne`, which streamlines the necessary corrections to raw drone data and produces operative thermal landscapes with minimal input and technical expertise required by the user. This approach should enhance the accuracy and accessibility of detailed and biologically relevant thermal landscapes for species around the globe and improve the quality of the baseline data that are required for forecasts of the responses of organisms to environmental change.

Our method works by linking temporally continuous (but spatially discrete) OTM thermal profiles to spatially continuous (but temporally discrete) thermal orthomosaics generated from drone photogrammetry to generate spatiotemporally comprehensive operative temperature maps (i.e., thermal landscapes). We confirmed that our approach can produce highly accurate thermal landscapes by comparing the predicted temperatures from the `throne` output to real operative temperatures measured with OTMs that we deployed in the field. Indeed, despite the temporal or spatial scale over which we made predictions, the average predictive error of final daytime thermal landscapes was < 0.5 °C, and 95% of the tile values within the landscapes were within 2 °C of the true value. This accuracy further increases when nighttime temperatures are included (as there is much less thermal heterogeneity at night). Nonetheless, the accuracy and thus utility of our method depends on the optimization of several factors.

First, successful implementation of this method depends on obtaining accurate *in-situ* operative temperature data using field-deployed data loggers (OTMs). In the field of thermal ecology, OTMs are designed in inconsistent ways and often are not properly calibrated (Dzialowski, 2005). However, recent advances in 3D printing provide opportunities to build affordable and accurate OTMs for a wide range of species (Alujević et al., 2024). Since our method involves correlating the thermal profiles of OTMs with tiles in the drone-acquired orthomosaics, it is crucial to deploy OTMs across diverse microhabitats, encompassing various substrates, vegetation types, and topographical features. This ensures a comprehensive range of options for linking OTM thermal profiles to tiles. The selection of microhabitats for OTM deployment should be informed by the ecology of the species under study. In our investigation, we strategically deployed OTMs with the goal of comprehensively sampling the microhabitats available to our study species. This allowed us to test the sensitivity of our method to OTM coverage in the thermally heterogeneous environment of the Great Basin Desert. Irrespective of whether we were predicting thermal landscapes across smaller or larger geographic areas, or over shorter or longer timeperiods, fewer OTMs than we expected were required to describe the thermal dynamics of the site. For an area of 33,000 m^2^ and over a 76-day period (validation 1), we found that only five OTMs were required to accurately describe half of the thermal variation at our site, while only 22 OTMs were required to accurately describe 90% of the thermal variation. This promising result suggests that, as long as OTMs are deployed strategically to capture key microhabitat and physiographic features of the site, researchers may not need to deploy large numbers of OTMs for optimal implementation of our method, especially in environments that are less thermally heterogenous than a rocky desert. Regardless, decisions on how many OTMs to deploy at a given site will be system specific.

A second consideration is the appropriate smoothing factor (i.e., the number of knots per hour; Figure S14) to generate daily thermal profiles for each OTM. This decision depends on two factors: the frequency at which OTMs recorded operative temperatures and the biophysical ecology of the study organism. While it is essential to capture accurate thermal profiles of specific microsites, excessive precision in these profiles will introduce noise that is irrelevant to the study organism and might negatively impact the quality of the match between the thermodynamics of OTMs and tiles from the drone data. What counts as “excessive precision” will be system-dependent. For example, in an environment where abrupt changes in weather (e.g., gusts of wind, brief cloud cover, etc.) result in rapid and reversible shifts in temperature that are unlikely to influence the behavior of the organism (because the particular organism has relatively high body mass and therefore relatively high thermal inertia, for example), smoothing improves the quality of the final estimated thermal landscape. This was generally the case for our focal population of the western fence lizard (average adult mass > 20 g), and we therefore opted to use 4 knots per hour which smoothed OTM measurements at 15-minute intervals. However, for smaller bodied organisms that are more susceptible to short-term changes in abiotic conditions (e.g., lizards or insects under 5 g in mass), one might increase the number of knots in order to capture shorter-term thermal fluctuations in the environment.

A third factor that we thought would influence the quality of the thermal landscapes produced by `throne` was the number of drone missions flown over the field site. Although we had assumed that a larger number of flights would be advantageous as it would provide more time points to correlate OTM thermal profiles with tile dynamics, our results show that a relatively small number of flights is probably sufficient for most applications. Regardless of whether thermal landscapes were generated using 3 randomly selected flights or 34, and whether operative temperature predictions were made for longer or shorter time periods, or over larger or smaller geographic areas, the accuracy of `throne`’s output was similar (Table 1). This result is likely robust as we tested our method in the highly heterogeneous environment of a high-elevation, temperate desert. This is a notable advantage of `throne`, as the researcher or practitioner will only have to fly their drone a moderate number of times to accurately capture the long-term thermal dynamics of their field site. This should increase the number of missions or size of the field site that can be measured under the same drone battery power, which can be an important rate-limiting factor in drone-based studies.

Finally, it is important to consider the spatial resolution of drone-based surface temperatures that is necessary to capture thermal variation at a scale that is relevant to the study organism. There are two parameters to consider: 1) the resolution that can be attained by the specific brand and model of TIR camera being used, and 2) the resolution of the resulting thermal landscapes that are produced with `throne` (Figure S15). TIR cameras typically produce images with a lower digital resolution than standard cameras (the Zenmuse XT2 thermal camera that we used has a resolution of 640 x 512 pixels). Flight altitude is also important to consider as it affects both the spatial resolution captured by the thermal camera (the size of the pixel on the ground represented by the GSD) and the amount of atmospheric interference between the ground and the sensor (Playà-Montmany & Tattersall 2021). Users should adjust the flight altitude to generate thermal orthomosaics with a GSD that will ultimately generate a map with a resolution relevant to their focal organism. For example, at a flight altitude of 40 m, the center distance between two adjacent pixels in our thermal photos was 5.23 cm (each pixel was 0.027 m^2^ in area). A flight conducted at lower altitude could be needed for small organisms such as insects. However, using drone temperature data with an extremely low GSD might introduce noise that reduces the predictive capacity of the model. The spatial resolution of the final thermal landscape can also be modulated in `throne` with the argument `resolution` in the `rnp_flights_data` function. Although it would seem logical to always maximize the resolution such that 1 tile represents the smallest possible area (0.5 m^2^), increased spatial resolution requires greater computing power and time (especially during the tile-to-OTM matching step), which may not be available to some users. Despite these considerations, our approach will allow researchers to generate thermal landscapes for their study organisms with substantially higher resolution than those commonly used in climate forecasts (e.g. the WorldClim 2 dataset; https://www.worldclim.org/; Fick & Hijmans, 2017).

Major benefits of the `throne` package are that the expertise required to model thermal landscapes at nearly unprecedented levels of spatiotemporal detail is relatively low, the equipment required is relatively affordable, and the thermal landscape model produced is based on real operative temperatures collected at the specific site of interest. Ecophysiological modeling software such as NicheMapR (Kearney & Porter, 2017), trenchR (Buckley et al., 2023), microclima (Maclean et al., 2019), and others are useful tools for microclimate modeling, not only in current but also past and future climates (Trew et al. 2024, MacLean 2020), something our method can also do, and at high spatial resolution, but not without applying key assumptions that should be independently validated. For example, one could forecast operative temperature distributions by projecting IPCC predictions for increases in mean temperature onto the spatial roster from the throne output. While ecophysiological models are invaluable for simulating body temperatures at broader geographic scales, our approach allows for spatially explicit, high-resolution mapping of operative temperatures that are relevant to individual organisms without the need for assumptions or separate measurements of microhabitat frequencies and distributions. Further, our method is applicable to a wide range of open and semi-open environments that occur over broad swaths of the planet.

We have designed `throne` to be user-friendly in that it requires minimal analytical expertise; users can simply input their raw data and a few boundary parameters, and `throne` will generate predicted thermal landscapes for any day/time combination during which OTMs were logging temperatures in the field. To generate the input data, some training and expertise is required, but we do not think this will be insurmountable for most researchers. Thermal biologists have been building and deploying OTMs for decades (Angilletta, 2009; Bakken, 1992; Dzialowski, 2005), and modern commercially available drones with TIR cameras are both relatively affordable and easy to operate. Drones equipped with thermal sensors are rapidly becoming more affordable; the model used in this study was priced at ∼$8,000 in 2019 and the thermal sensor had to be purchased separately (∼$13,000). Only a few years later, drones with similar capabilities and built-in with thermal sensors have become much smaller and more portable, with options like the DJI Mavic 3 Thermal now available for as little as $5,000. Although the photogrammetry software used in this study (Pix4D) requires a license, there are several equivalent products that are now available for free (e.g., OpenDroneMap online software: https://www.opendronemap.org/). Despite this, some researchers (especially some of those in developing nations) may find our method too expensive or impractical, perhaps until drone and sensor prices decline further. Regardless, we see the accessibility of our method as a significant advance in the field because it will enable many researchers, practitioners, or conservation managers to precisely characterize the thermal environment of interest in a way that is both spatiotemporally comprehensive and relevant to the biology of their study organism.

Although our approach offers numerous benefits, there will be contexts in which it may not be the most appropriate method for quantifying operative thermal landscapes. First, it relies on *in situ* operative temperature measurements to convert drone-based surface temperatures to temperatures that are biologically appropriate for the study species. This means that `throne` can only be applied during the time period that operative temperatures are being logged by field deployed OTMs. In this study, we used OTMs that had Thermocron iButton temperature loggers suspended inside. However, even high-capacity iButtons can only record a few thousand temperature measurements. To create thermal landscapes for many months or the entire year, we would have had to retrieve the temperature loggers, download the data, and redeploy them several times. This process might not be feasible in other systems where accessing the OTMs is challenging or data storage capacity is limited. Finally, our approach is ideal for environments that are at least moderately open (e.g., deserts, grasslands, open woodlands, etc.). This is because drone thermal imagery can, by its nature, only capture temperatures of surfaces below the flight path. In habitats like rainforests with fully enclosed canopy, this approach would not be suitable as the drone sensors would be unlikely to penetrate to the understory except in cases where there were canopy gaps (but see alternative approach in Higgins et al., 2024). Nonetheless, our drone-based approach to mapping thermal environments is probably unnecessary in these types of highly buffered, thermally homogeneous environments as the deployment of a small number of temperature loggers usually captures the operative thermal dynamics in these habitats (Cox, Alexander, et al., 2020; Nicholson et al., 2022; Williams et al., 2022). Furthermore, in addition to its primary application for high-resolution operative thermal mapping, throne could also be implemented using .tif format datasets from satellite-derived surface temperature products, such as those from VIIRS, Copernicus, or MODIS, which offer coarser spatial resolution but can cover broader geographic regions. While this may not capture fine-scale thermal variability, it could still provide valuable insights for large-area studies when combined with ground-based OTMs.

The accurate mapping of terrestrial thermal environments has direct implications for predicting organismal responses to rapid environmental change. Inaccurate or low-resolution estimates of operative temperature can reduce our ability to quantify constraints on thermoregulatory capacity which generates inaccuracies and biases in our predictions for how organisms might respond to changing environments. Our drone-based framework for quantifying operative thermal environments offers a straightforward approach that produces accurate and precise spatiotemporal operative temperature distributions. The broad adoption of this approach should deepen our understanding of how organisms interact with their environments, how they have adapted to these environments in the past, and how climate change is likely to impact ecosystems in the future.

## Acknowledgements

We would like to thank Matthew Gifford, Christopher Halsch, Matt Forister, Lee Dyer, Madeleine Lohman and Julia Brockman for their helpful feedback while developing this method and, especially, Matthieu Bruneaux for the thorough review and feedback provided on the R package.

## Author Contributions

K.A., G.G-C., J.B. and M.L.L. conceived the ideas and designed methodology; K.A., G.G.-C., N.R. and A.C.G. collected the data; K.A., G.G.-C., N.R., E.S. and R.G. analysed the data; G.G.-C led the development of the R package; K.A. and G.G.-C. led the writing of the manuscript. All authors contributed critically to the drafts and gave final approval for publication.

