## Supplementary material for "Using aerial thermography to map terrestrial thermal environments in unprecedented detail"

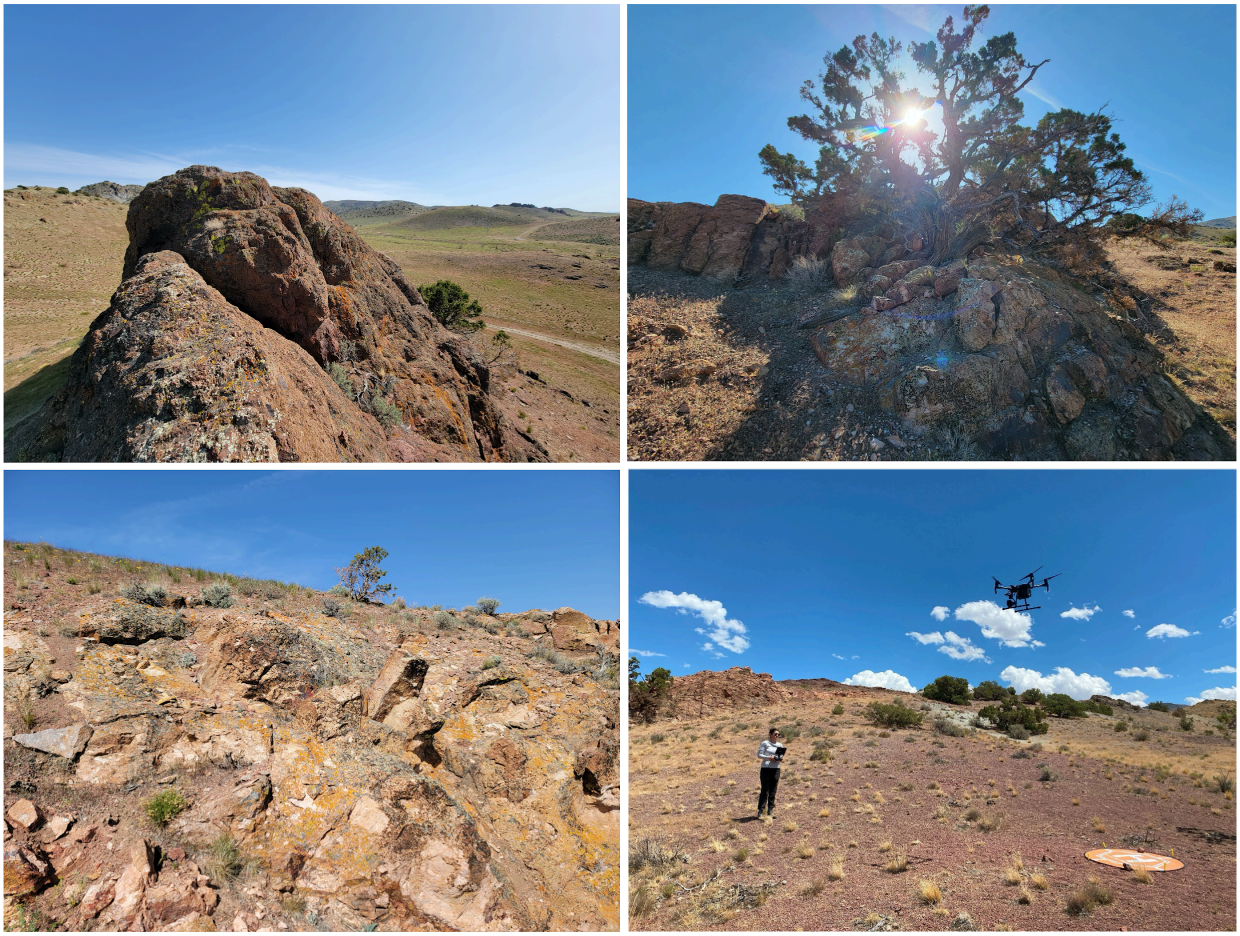

**Figure S1.** Our study site in the Great Basin Desert.

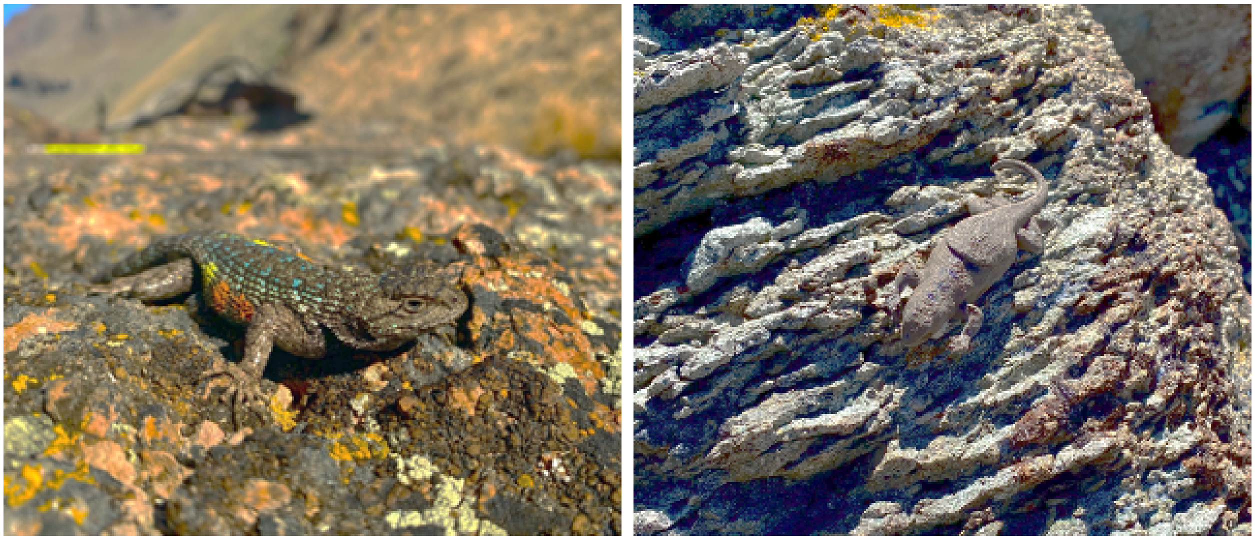

**Figure S2.** Western fence lizard (*Sceloporus occidentalis*) on the left and OTM on the right.

**Table S1**. List of microhabitats where OTMs were deployed with corresponding descriptions and numbers of OTMs deployed (N) for validation 1 and 2.

| Validation 1 | | |
|---|---|---|
| Microhabitat | Description | N |
| Rock N | Large, isolated *or* connected rock surface, predominantly facing north* | 8 |
| Rock E | Large, isolated *or* connected rock surface, predominantly facing east* | 8 |
| Rock S | Large, isolated *or* connected rock surface, predominantly facing south* | 9 |
| Rock W | Large, isolated *or* connected rock surface, predominantly facing west* | 7 |
| Ground | Gravel or coarse sand substrate | 9 |
| Bush | Great Basin sagebrush (*Artemisia tridentata*) bush | 9 |
| Tree | Utah juniper (*Juniperus osteosperma*) tree | 9 |
| Free | Freely placed within the study site, either category | 18 |
| Validation 2 | | |
| Outcrop | Large, exposed and connected rock surface, extending for at least 2 meters | 12 |

| | | |
|---|---|---|
| Boulder | Large, isolated rock on ground, between 1-2 m in diameter | 6 |
| Rock | Isolated rock on ground, < 1m in diameter | 8 |
| Ground | Gravel or coarse sand substrate | 9 |
| Bush | Great Basin sagebrush (Artemisia tridentata) bush | 1 |

*If the incline of the substrate was less than 25° (we arbitrarily chose this cutoff to account for exposure to sunlight) in all directions, the orientation was noted as "flat"; if the incline was greater than 25°, the orientation was assigned as one of cardinal directions: N, W, S, E.

**Table S2.** Flight metadata for validation 1 and 2.

| Validation | Date | Start time | Flight duration | Ambient temperature (°C) | Wind speed (m/s) | Flight altitude (m) | Cloud cover |
|---|---|---|---|---|---|---|---|
| 1 | 05/15/2023 | 18:13 | 12 | 24.61 | 3.75 | 100 | sunny/very few clouds |
| 1 | 05/16/2023 | 10:05 | 17 | 20.33 | 3.35 | 100 | sunny/no clouds |
| 1 | 05/18/2023 | 8:16 | 10 | 16.89 | 1.7 | 100 | sunny/no clouds |
| 1 | 06/02/2023 | 10:35 | 8 | 19.67 | 1.7 | 100 | sunny/no clouds |
| 1 | 06/02/2023 | 15:39 | 9 | 24.83 | 1.97 | 100 | sunny/no clouds |
| 1 | 06/19/2023 | 14:25 | 8 | 15.72 | 6.08 | 100 | sunny/very few clouds |
| 1 | 06/19/2023 | 16:55 | 9 | 18.17 | 5.77 | 100 | sunny/very few clouds |
| 1 | 07/05/2023 | 19:42 | 9 | 30 | 3.04 | 100 | mostly sunny/few clouds |
| 1 | 07/09/2023 | 12:35 | 8 | 32.28 | 0.31 | 100 | sunny/no clouds |
| 1 | 07/29/2023 | 11:00 | 9 | 32.89 | N/A | 100 | mostly sunny/few clouds |
| 2 | 08/24/2023 | 8:35 | 4 | 18.67 | 4.69 | 40 | sunny/no clouds |
| 2 | 08/24/2023 | 8:59 | 4 | 21.33 | 3.67 | 40 | sunny/no clouds |
| 2 | 08/24/2023 | 9:19 | 4 | 21.94 | 2.91 | 40 | sunny/no clouds |
| 2 | 08/24/2023 | 9:42 | 4 | 23.56 | 3.8 | 40 | sunny/no clouds |
| 2 | 08/24/2023 | 10:06 | 4 | 24.39 | 3.04 | 40 | sunny/no clouds |
| 2 | 08/24/2023 | 10:38 | 4 | 24.89 | 2.99 | 40 | sunny/no clouds |
| 2 | 08/24/2023 | 10:57 | 4 | 26 | 2.46 | 40 | sunny/no clouds |

| | | | | | | | |
|---|---|---|---|---|---|---|---|
| 2 | 08/24/2023 | 11:20 | 4 | 26.67 | 4.25 | 40 | sunny/no clouds |
| 2 | 08/24/2023 | 11:40 | 4 | 28.06 | 4.96 | 40 | sunny/very few clouds |
| 2 | 08/24/2023 | 12:00 | 4 | 26.78 | 2.46 | 40 | sunny/very few clouds |
| 2 | 08/24/2023 | 12:21 | 4 | 26.33 | 4.47 | 40 | mostly cloudy |
| 2 | 08/25/2023 | 10:27 | 4 | 25.28 | 4.69 | 40 | sunny/no clouds |
| 2 | 08/25/2023 | 10:49 | 4 | 25.56 | 3.22 | 40 | sunny/no clouds |
| 2 | 08/25/2023 | 11:09 | 4 | 26.39 | 4.69 | 40 | sunny/no clouds |
| 2 | 08/25/2023 | 11:30 | 4 | 26.22 | 4.6 | 40 | sunny/very few clouds |
| 2 | 08/25/2023 | 11:52 | 4 | 27.33 | 4.38 | 40 | sunny/some clouds |
| 2 | 08/25/2023 | 12:12 | 4 | 27.17 | 4.43 | 40 | sunny/some clouds |
| 2 | 08/25/2023 | 12:37 | 4 | 27.39 | 4.83 | 40 | sunny/some clouds |
| 2 | 08/25/2023 | 12:57 | 4 | 28.33 | 3.31 | 40 | mostly cloudy |
| 2 | 08/25/2023 | 13:18 | 4 | 28.28 | 4.02 | 40 | mostly cloudy |
| 2 | 08/25/2023 | 13:39 | 4 | 29.94 | 3.84 | 40 | sunny/some clouds |
| 2 | 08/25/2023 | 14:00 | 4 | 29.06 | 3.62 | 40 | mostly cloudy |
| 2 | 08/26/2023 | 14:36 | 4 | 29.67 | 5.86 | 40 | sunny/no clouds |
| 2 | 08/26/2023 | 15:00 | 4 | 29.72 | 3.75 | 40 | sunny/no clouds |
| 2 | 08/26/2023 | 15:26 | 4 | 29.78 | 5.19 | 40 | sunny/very few clouds |
| 2 | 08/26/2023 | 15:51 | 4 | 29.11 | 3.8 | 40 | sunny/very few clouds |
| 2 | 08/26/2023 | 16:16 | 4 | 29.72 | 6.03 | 40 | sunny/some clouds |
| 2 | 08/26/2023 | 16:38 | 4 | 30.39 | 6.17 | 40 | sunny/some clouds |
| 2 | 08/26/2023 | 17:06 | 4 | 29.44 | 3.8 | 40 | sunny/some clouds |
| 2 | 08/26/2023 | 17:31 | 4 | 30.06 | 2.19 | 40 | sunny/very few clouds |
| 2 | 08/26/2023 | 17:56 | 4 | 29.33 | 3.31 | 40 | sunny/very few clouds |
| 2 | 08/26/2023 | 18:21 | 4 | 29.33 | 0.72 | 40 | sunny/no clouds |
| 2 | 08/26/2023 | 18:46 | 4 | 30.22 | 1.56 | 40 | sunny/no clouds |

| | | | | | | | |
|---|---|---|---|---|---|---|---|
| 2 | 08/26/2023 | 19:08 | 4 | 29.67 | 0 | 40 | sunny/no clouds |

**Table S3.** Mean predictive error (± SD) for validation 1 and 2, presented as the absolute difference between the observed (from OTMs) and predicted (from the final thermal landscape output by throne) temperatures across different combinations of number of drone flights, number of OTMs, and knot_p values (magnitude of smoothing of raw OTM data; knots per hours are given in the brackets) used to generate the final thermal landscape. All comparisons were done for data collected during daytime hours (7 AM to 7 PM).

| **Validation 1** | | **knot_p (knot/h)** | | | |
|---|---|---|---|---|---|
| **N Flight** | **N OTM** | **0.25 (~0.3)** | **0.5 (~0.6)** | **1 (~1.2)** | |
| **3** | 20 | 0.575 ± 0.369 | 0.469 ± 0.282 | 0.388 ± 0.411 | |
| | 30 | 0.46 ± 0.368 | 0.401 ± 0.255 | 0.242 ± 0.2 | |
| | 70 | 0.474 ± 0.262 | 0.221 ± 0.249 | 0.792 ± 0.388 | |
| **5** | 20 | 0.956 ± 0.669 | 0.392 ± 0.261 | 0.666 ± 0.292 | |
| | 30 | 0.879 ± 0.611 | 0.396 ± 0.162 | 0.253 ± 0.154 | |
| | 70 | 0.909 ± 0.824 | 0.317 ± 0.244 | 0.29 ± 0.207 | |
| **10** | 20 | 0.64 ± 0.687 | 0.419 ± 0.442 | 0.315 ± 0.278 | |
| | **30** | 0.544 ± 0.296 | 0.375 ± 0.32 | 0.658 ± 0.496 | |
| | **70** | 1.152 ± 0.758 | 1.031 ± 0.656 | 0.46 ± 0.382 | |
| **Validation 2** | | **knot_p (knot/h)** | | | |
| **N Flight** | **N OTM** | **0.017 (0.5)** | **0.067 (2)** | **0.133 (4)** | **0.5 (15)** |
| 3 | 10 | 0.911 ± 0.435 | 0.85 ± 0.618 | 0.828 ± 0.275 | 0.743 ± 0.506 |
| | 20 | 0.447 ± 0.259 | 0.729 ± 0.618 | 0.857 ± 0.415 | 0.577 ± 0.199 |

| | | | | | |
|---|---|---|---|---|---|
| | 33 | 0.554 ± 0.511 | 1.095 ± 0.541 | 0.898 ± 0.592 | 0.667 ± 0.233 |
| 8 | 10 | 0.697 ± 0.421 | 1.12 ± 0.624 | 0.66 ± 0.407 | 0.589 ± 0.223 |
| | 20 | 1.025 ± 0.602 | 0.971 ± 0.569 | 0.415 ± 0.191 | 0.633 ± 0.325 |
| | 33 | 0.961 ± 0.563 | 0.977 ± 0.625 | 0.945 ± 0.682 | 0.899 ± 0.565 |
| 17 | 10 | 0.848 ± 0.553 | 0.651 ± 0.502 | 0.624 ± 0.371 | 0.844 ± 0.459 |
| | 20 | 0.937 ± 0.483 | 0.634 ± 0.435 | 0.826 ± 0.574 | 0.807 ± 0.431 |
| | 33 | 1.02 ± 0.587 | 0.751 ± 0.426 | 0.791 ± 0.314 | 0.959 ± 0.441 |
| 34 | 10 | 0.983 ± 0.479 | 1.028 ± 0.578 | 0.819 ± 0.494 | 0.672 ± 0.484 |
| | 20 | 0.871 ± 0.513 | 0.827 ± 0.404 | 0.964 ± 0.441 | 0.872 ± 0.549 |
| | 33 | 0.873 ± 0.457 | 0.842 ± 0.469 | 0.869 ± 0.345 | 0.967 ± 0.503 |

**Table S4.** Mean predictive error (± SD) for validation 1 and 2, presented as the relative difference between the observed (from OTMs) and predicted (from the final thermal landscape output by `throne`) temperatures across different combinations of number of drone flights, number of OTMs, and `knot_p` values (magnitude of smoothing of raw OTM data; knots per hours are given in the brackets) used to generate the final thermal landscape. All comparisons were done for data collected across all hours of the day.

| **Validation 1** | | **`knot_p` (knot/h)** | | |
|---|---|---|---|---|
| **N Flight** | **N OTM** | **0.25 (~0.3)** | **0.5 (~0.6)** | **1 (~1.2)** |
| 3 | 20 | 0.132 ± 0.51 | 0.092 ± 0.389 | -0.165 ± 0.364 |
| | 30 | 0.218 ± 0.341 | 0.171 ± 0.285 | -0.138 ± 0.215 |
| | 70 | 0.227 ± 0.297 | 0.086 ± 0.217 | 0.276 ± 0.566 |
| 5 | 20 | 0.425 ± 0.696 | 0.159 ± 0.287 | -0.274 ± 0.428 |
| | 30 | 0.353 ± 0.65 | 0.041 ± 0.309 | -0.056 ± 0.204 |

| | | | | | |
|---|---|---|---|---|---|
| | 70 | 0.501 ± 0.68 | -0.05 ± 0.28 | -0.15 ± 0.2 | |
| 10 | 20 | 0.171 ± 0.639 | 0.149 ± 0.397 | 0.021 ± 0.308 | |
| | 30 | 0.183 ± 0.396 | -0.167 ± 0.302 | 0.272 ± 0.504 | |
| | 70 | 0.282 ± 0.917 | 0.285 ± 0.805 | 0.136 ± 0.397 | |
| **Validation 2** | | `knot_p` **(knot/h)** | | | |
| **N Flight** | **N OTM** | **0.017 (0.5)** | **0.067 (2)** | **0.133 (4)** | **0.5 (15)** |
| 3 | 10 | 0.108 ± 0.729 | -0.263 ± 0.711 | 0.037 ± 0.629 | 0.102 ± 0.692 |
| | 20 | -0.116 ± 0.432 | -0.375 ± 0.581 | 0.043 ± 0.707 | -0.03 ± 0.511 |
| | 33 | -0.233 ± 0.49 | -0.1 ± 0.894 | -0.018 ± 0.831 | -0.008 ± 0.603 |
| 8 | 10 | 0.001 ± 0.664 | 0.097 ± 0.96 | 0.018 ± 0.654 | -0.101 ± 0.611 |
| | 20 | 0.014 ± 0.936 | 0.043 ± 0.857 | -0.135 ± 0.436 | -0.169 ± 0.61 |
| | 33 | -0.227 ± 0.847 | -0.203 ± 0.857 | -0.19 ± 0.854 | -0.188 ± 0.826 |
| 17 | 10 | -0.04 ± 0.783 | 0.041 ± 0.628 | -0.124 ± 0.585 | 0.122 ± 0.72 |
| | 20 | -0.146 ± 0.809 | -0.125 ± 0.642 | -0.001 ± 0.794 | -0.079 ± 0.737 |
| | 33 | -0.277 ± 0.883 | -0.206 ± 0.721 | -0.185 ± 0.715 | -0.161 ± 0.889 |
| 34 | 10 | -0.007 ± 0.831 | 0.175 ± 0.834 | 0.159 ± 0.699 | 0.133 ± 0.588 |
| | 20 | -0.071 ± 0.788 | -0.071 ± 0.747 | -0.152 ± 0.85 | -0.039 ± 0.838 |
| | 33 | -0.216 ± 0.79 | -0.2 ± 0.797 | -0.215 ± 0.836 | -0.103 ± 0.926 |

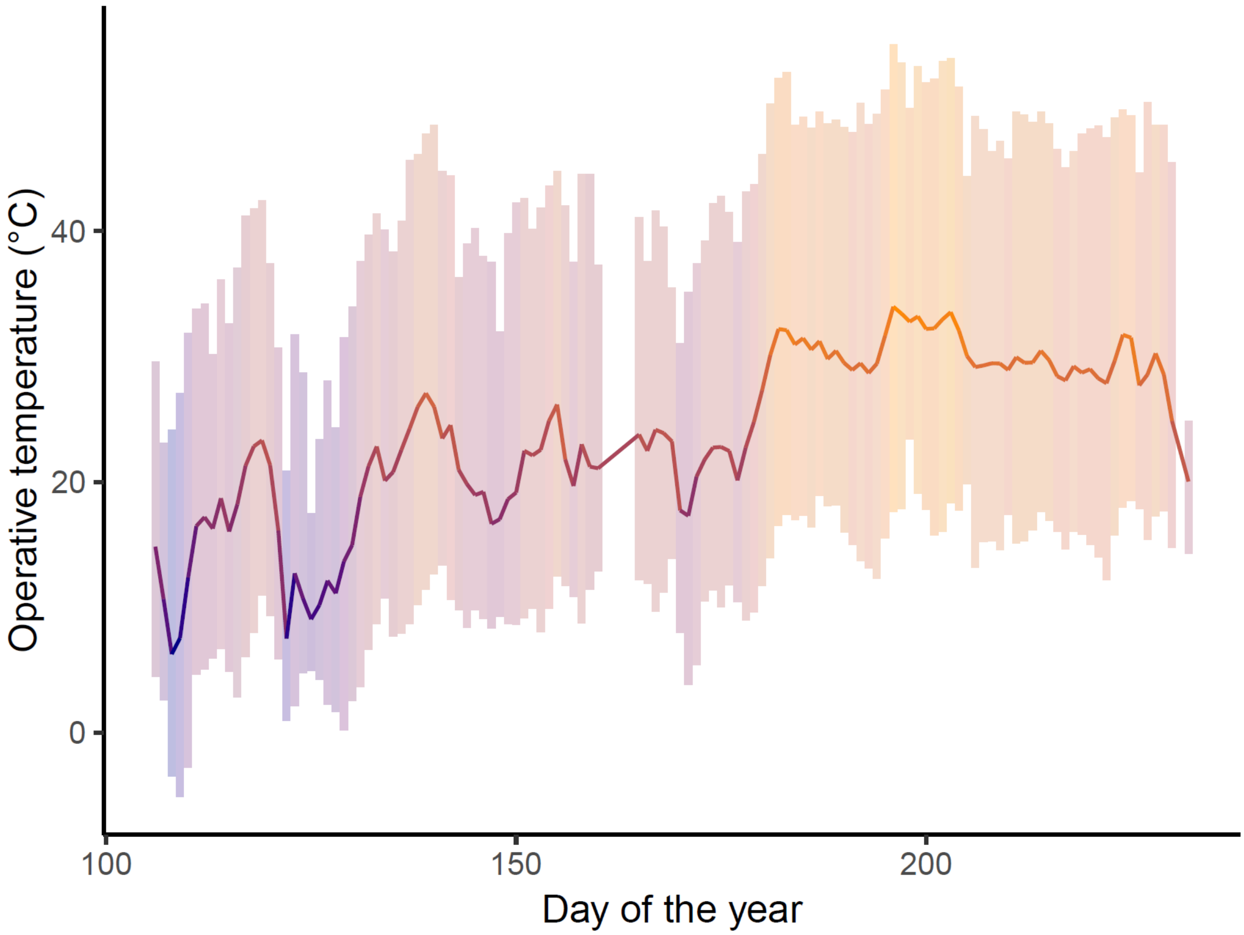

**Figure S3.** Fluctuation in operative temperature over the validation 1 period. Line represents mean and error bars show daily average minimum and maximum temperatures across all OTMs. The gap with no data corresponds to the 3-day period when OTMs were retrieved from the field, processed to obtain data and re-deployed.

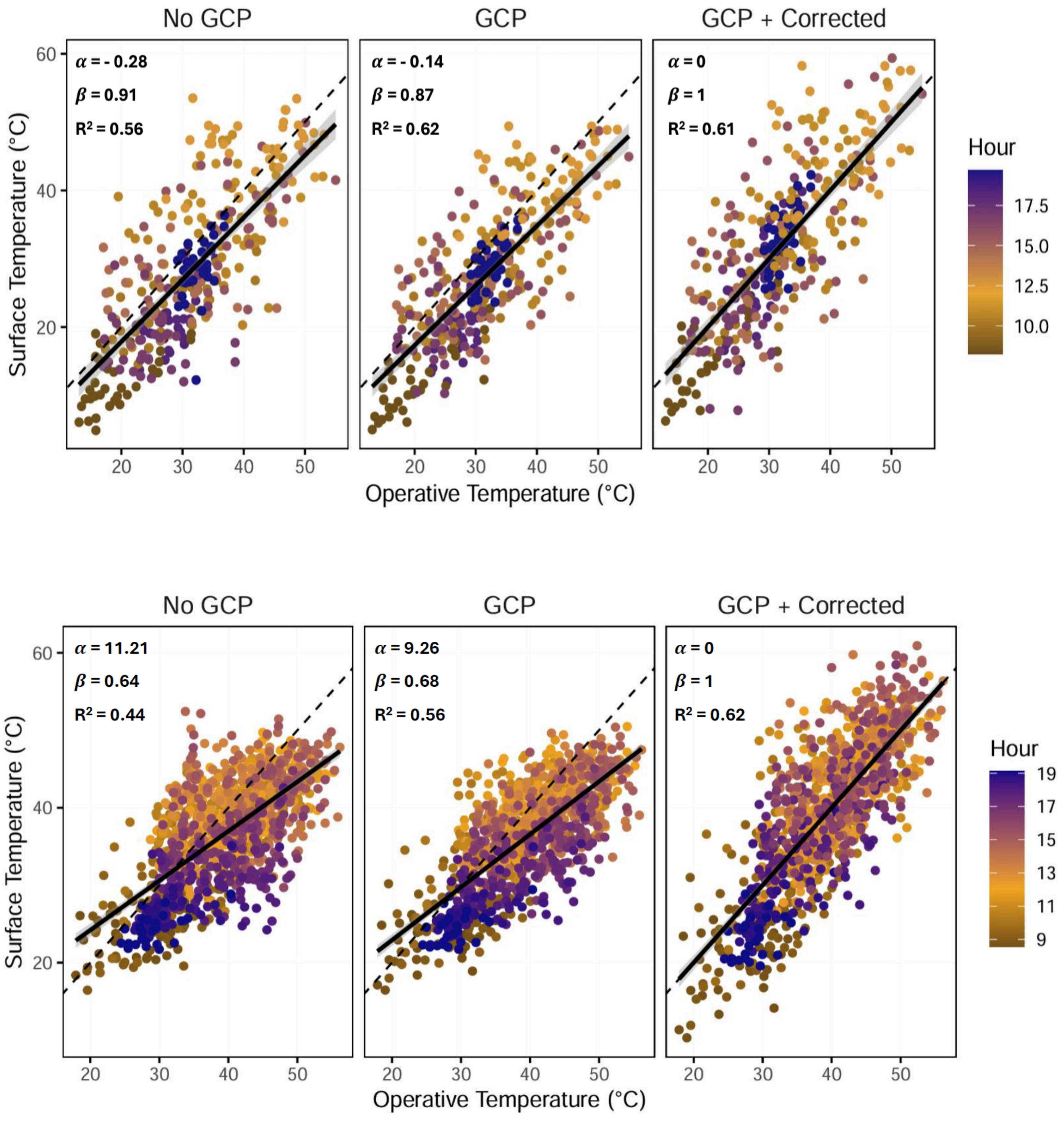

**Figure S4.** Correlation between operative temperatures and surface temperatures before applying corrections (left panel), after correcting for GCPs (middle panel), and after correcting for GCPs and the `correct_flights_data` function of the `throne` package (right panel) for validation 1 (top) and validation 2 (bottom).

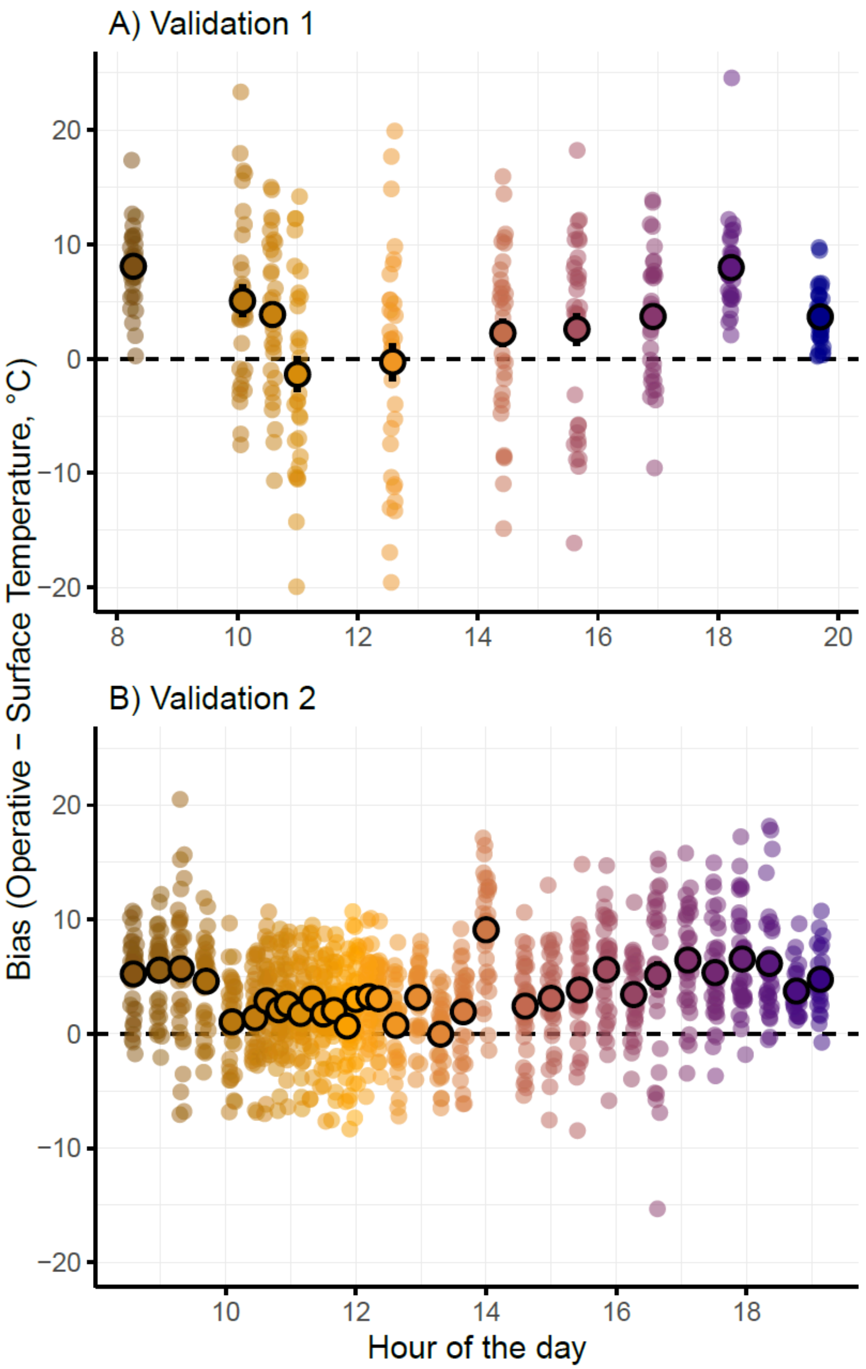

**Figure S5.** Temperature bias observed for validation 1 (A) and validation 2 (B) expressed as a difference between operative and drone surface temperatures as a function of time of day. Dots outlined in black represent means and bars represent standard errors.

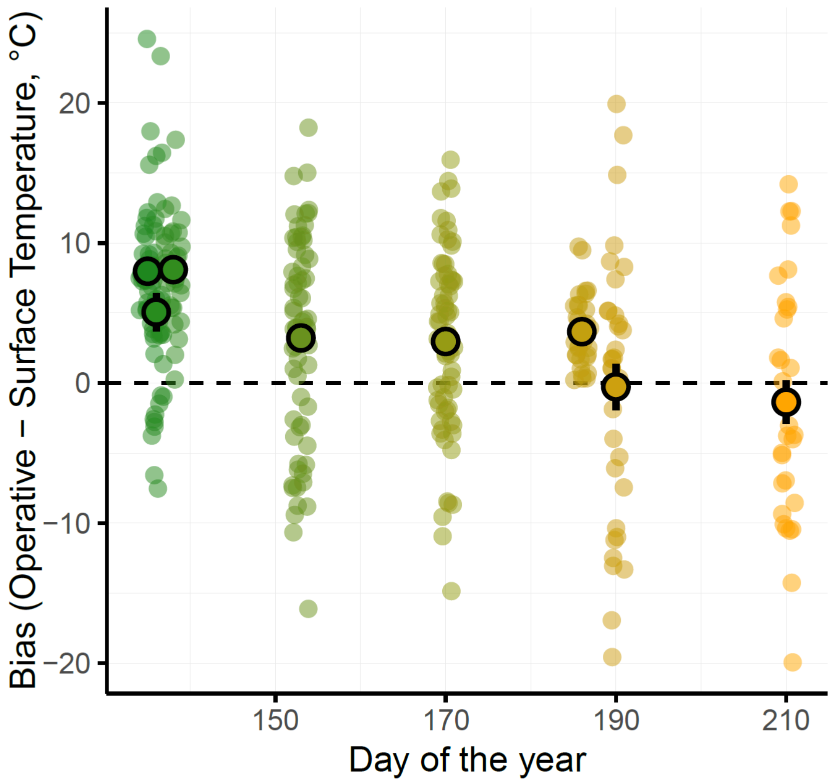

**Figure S6.** Temperature bias observed for validation 1 is expressed as a difference between operative and drone surface temperatures as a function of day of the year. Dots outlined in black represent means and bars represent standard errors.

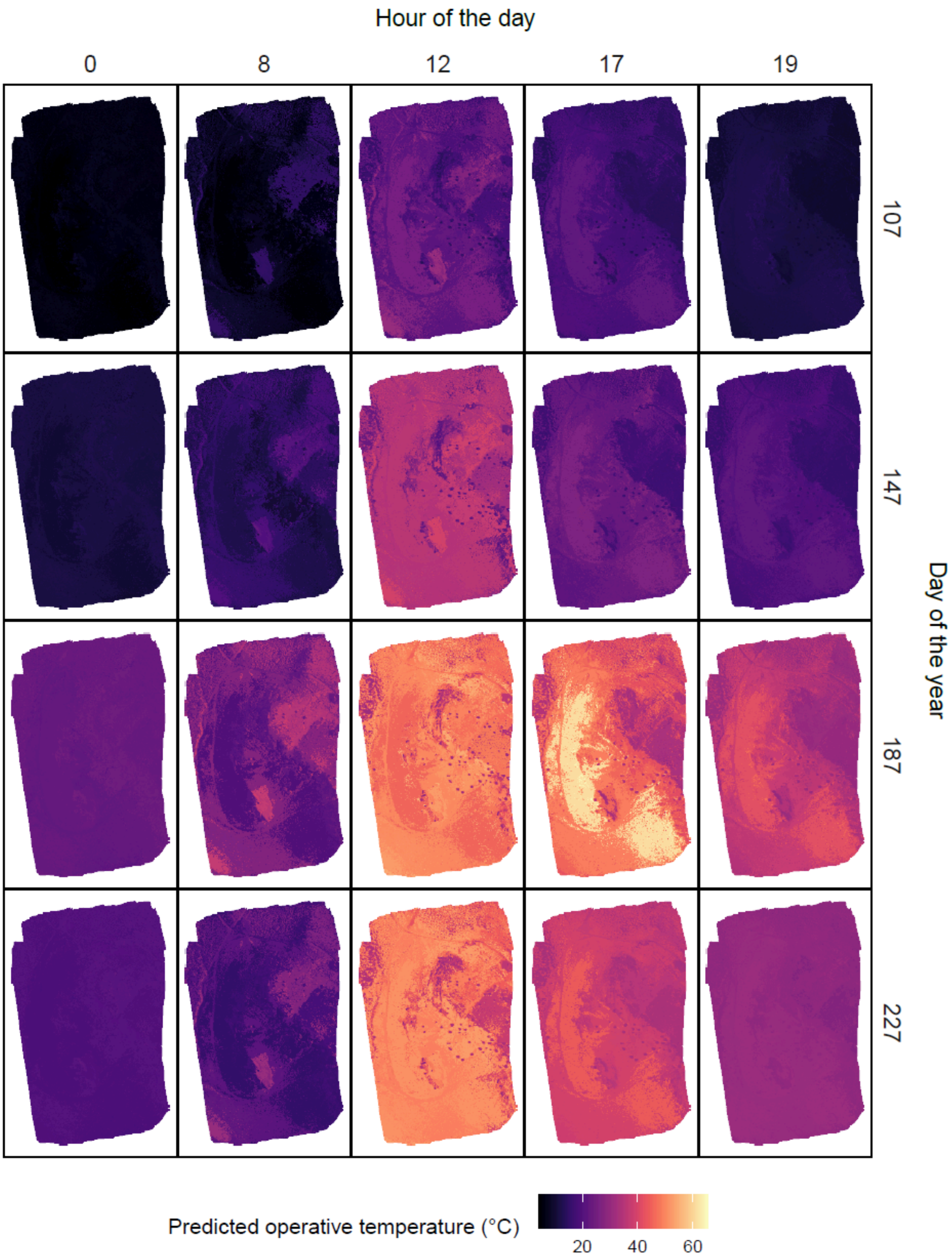

Hour of the day
0
8
12
17
19
107
147
187
227
Day of the year
Predicted operative temperature (°C)
20
40
60

**Figure S7.** Predicted thermal landscape generated for our study site in Northern Nevada generated for four Julian dates at five time points of the day..

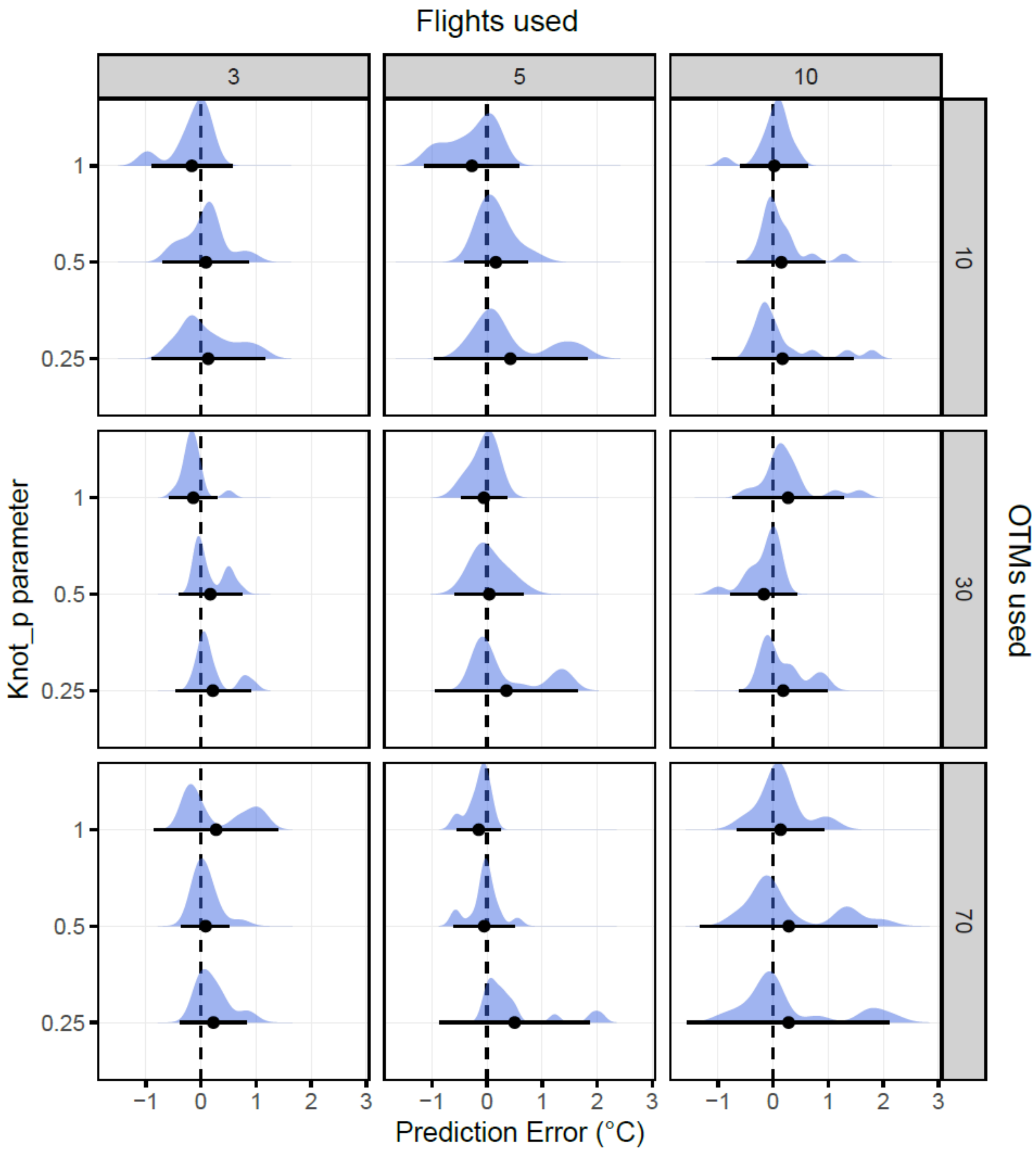

**Figure S8.** Daily frequency distribution of error (predicted-observed temperature) across different combinations of `knot_p` parameters, numbers of OTMs and drone flights for validation 1. Dots represent mean values and bars represent a 95% confidence interval around the mean. Note that the `knot_p` parameters are equivalent to $1 \sim 1.2$ knot/h , $0.5 \sim 0.6$ knot/h,

0.25 ~ 0.3 knot/h.

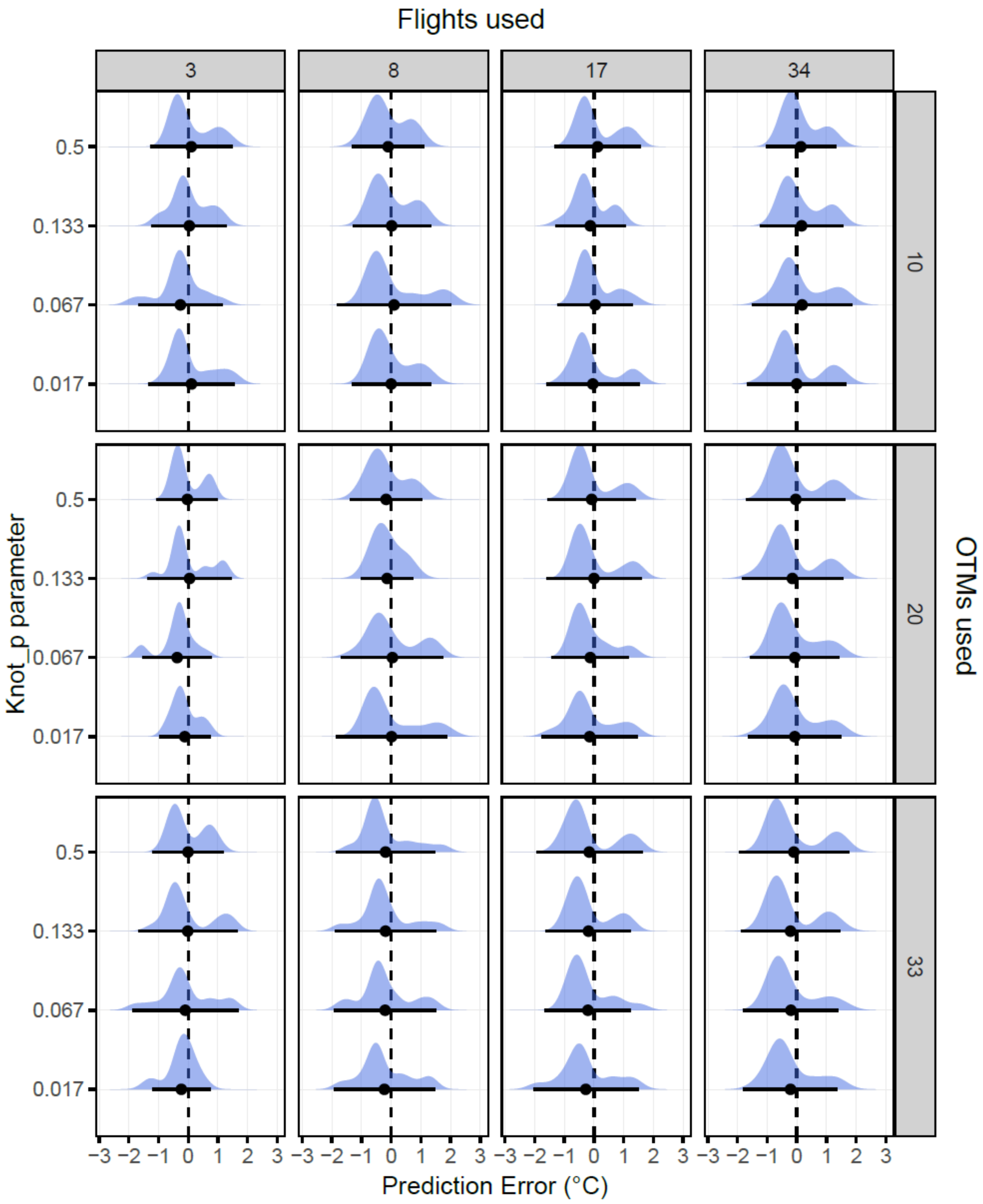

Flights used
3
8
17
34
10
20
33
OTMs used
Knot_p parameter
0.5
0.133
0.067
0.017
−3 −2 −1 0 1 2 3
Prediction Error (°C)

**Figure S9.** Daily frequency distribution of error (predicted-observed temperature) across different combinations of `knot_p` parameters, numbers of OTMs and drone flights for validation 2.

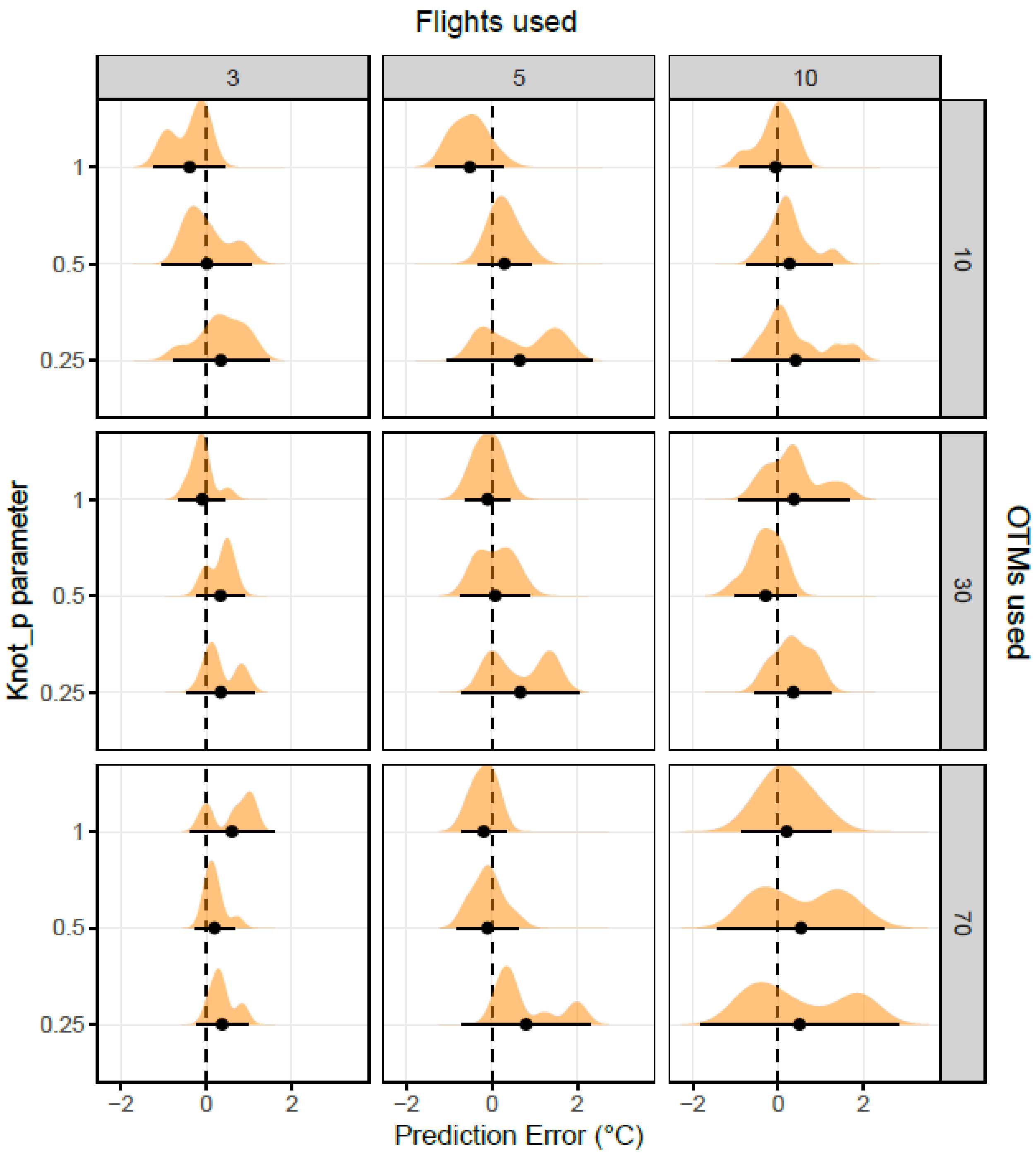

**Figure S10.** Daytime only frequency distribution of error (predicted-observed temperature) across different combinations `knot_p` parameters, numbers of OTMs and drone flights for validation 1. Dots represent mean values and bars represent 95% confidence intervals around the mean. Note that the `knot_p` parameters are equivalent to 1 ~ 1.2 knot / h, 0.5 ~ 0.6 knot/h, 0.25 ~ 0.3 knot/h,

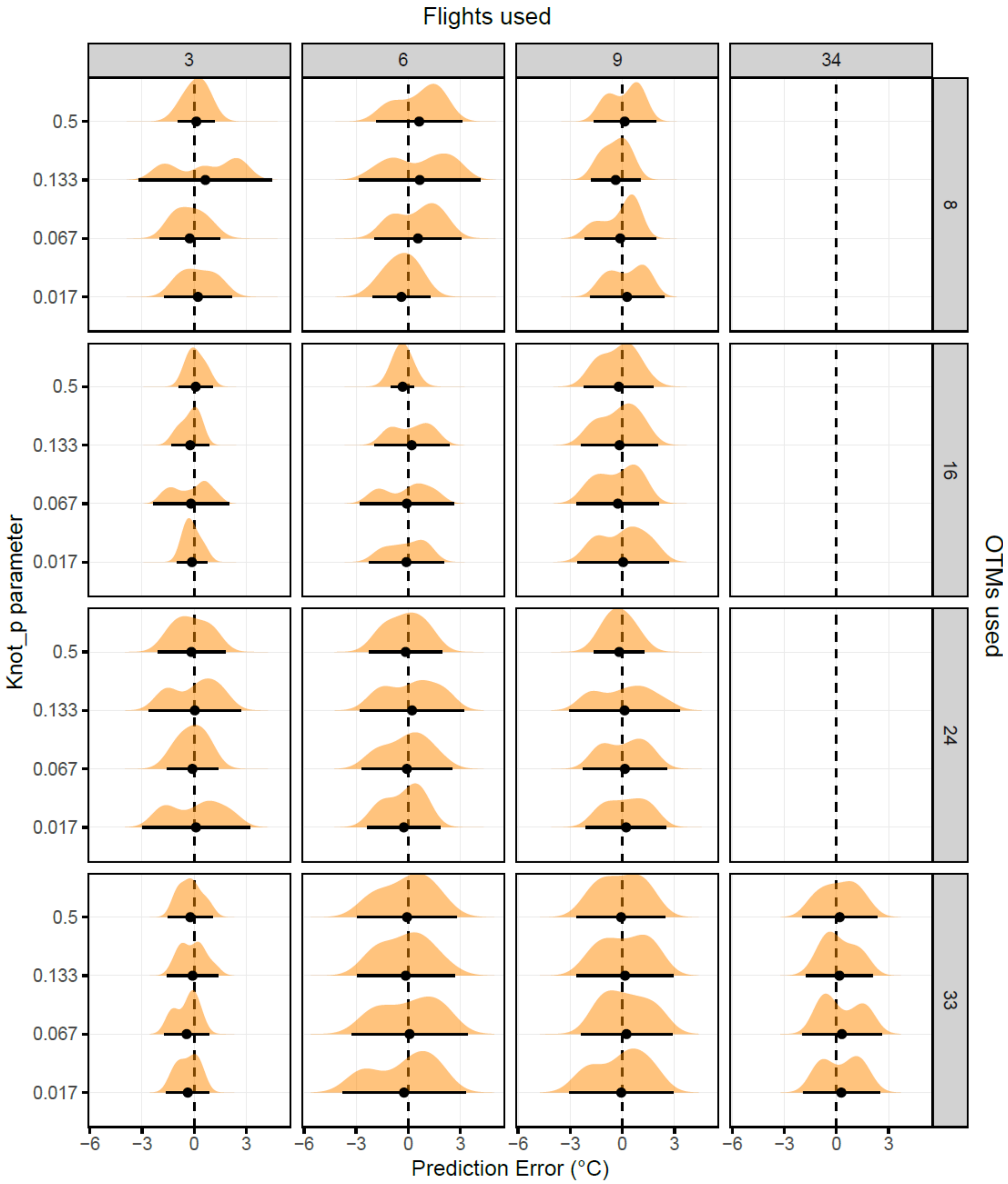

**Figure S11.** Daytime only frequency distribution of error (predicted-observed temperature) across different combinations `knot_p` parameters, numbers of OTMs and drone flights for validation 1.

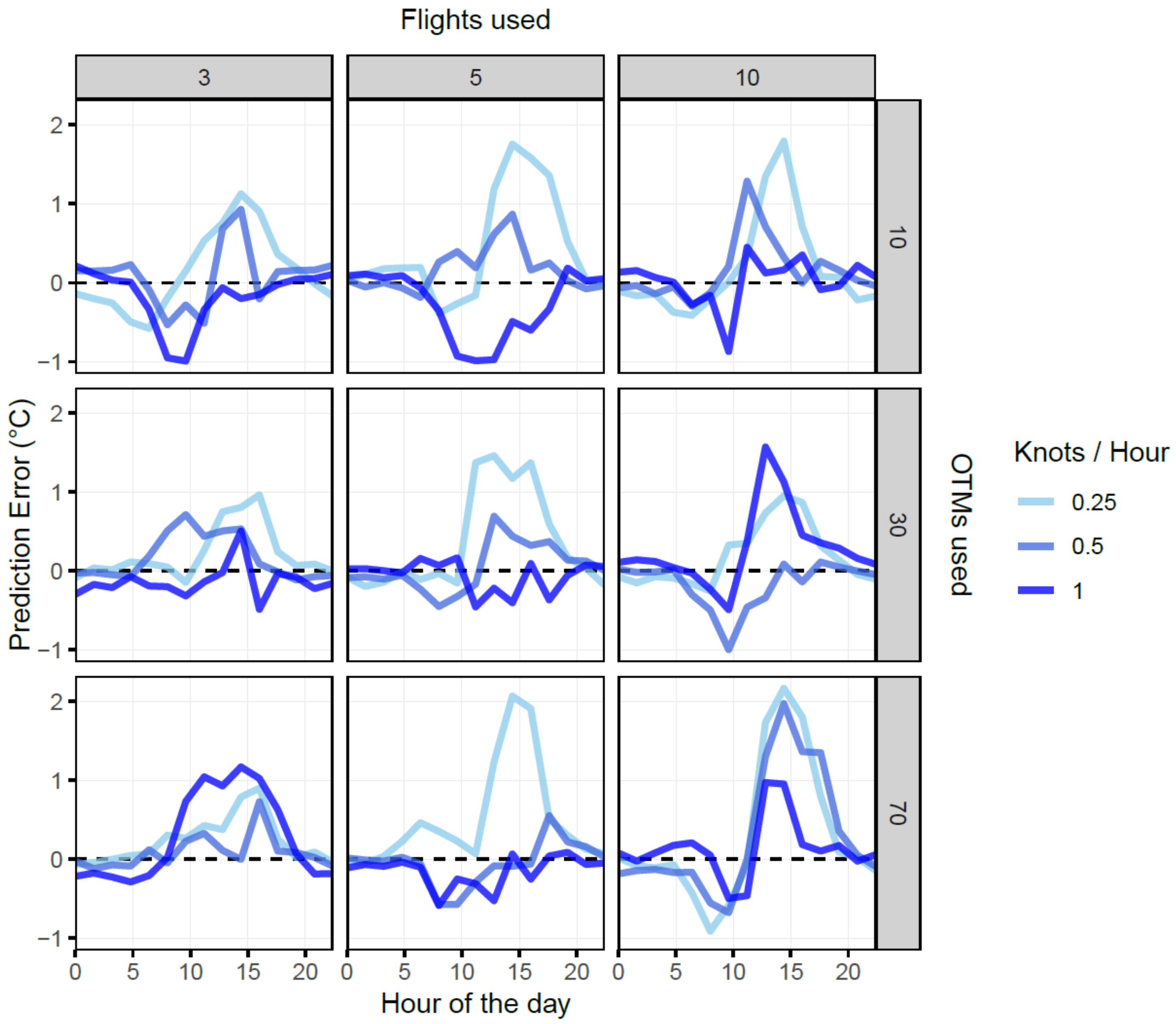

**Figure S12.** Prediction error as a function of time of day across different combinations of `knot_p` parameters, numbers of OTMs and drone flights for validation 1. Note that the `knot_p` parameters are equivalent to 1 ~ 1.2 knot/h, 0.5 ~ 0.6 knot/h, and 0.25 ~ 0.3 knot/h.

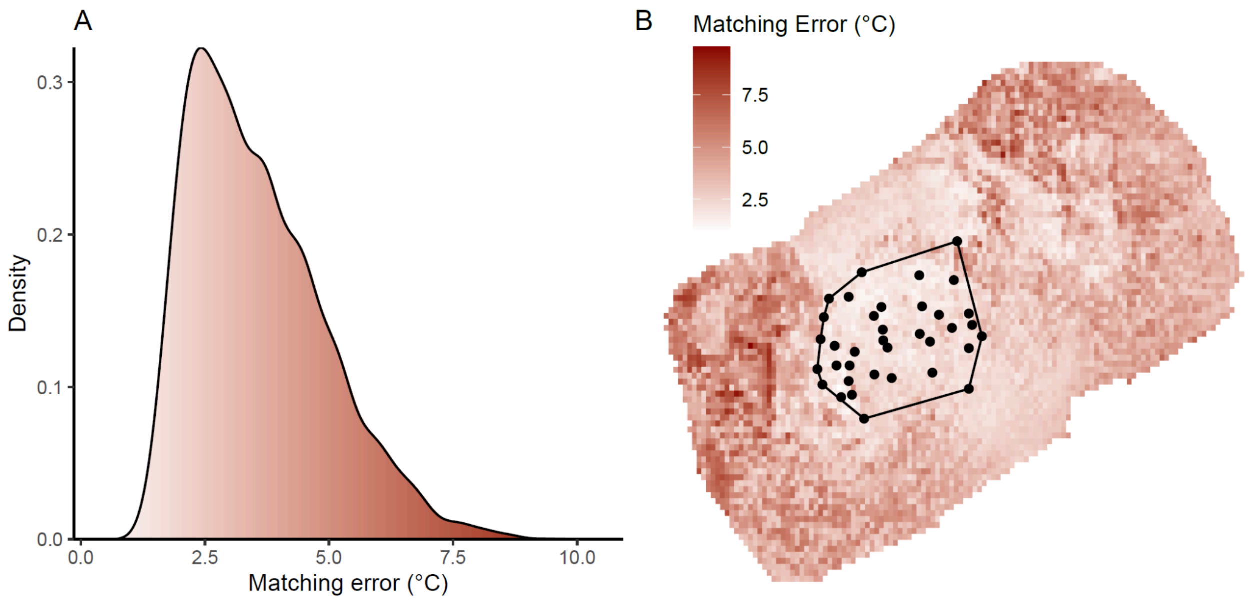

**Figure S13.** Matching error between tiles and the OTMs that best describe them for validation 2. Most of the error falls below 5 °C, the limit we recommend (A), further, the areas with the greater error are areas that surround the area of interest (highlighted by the positions of the OTMs deployed in the ground in black, B).

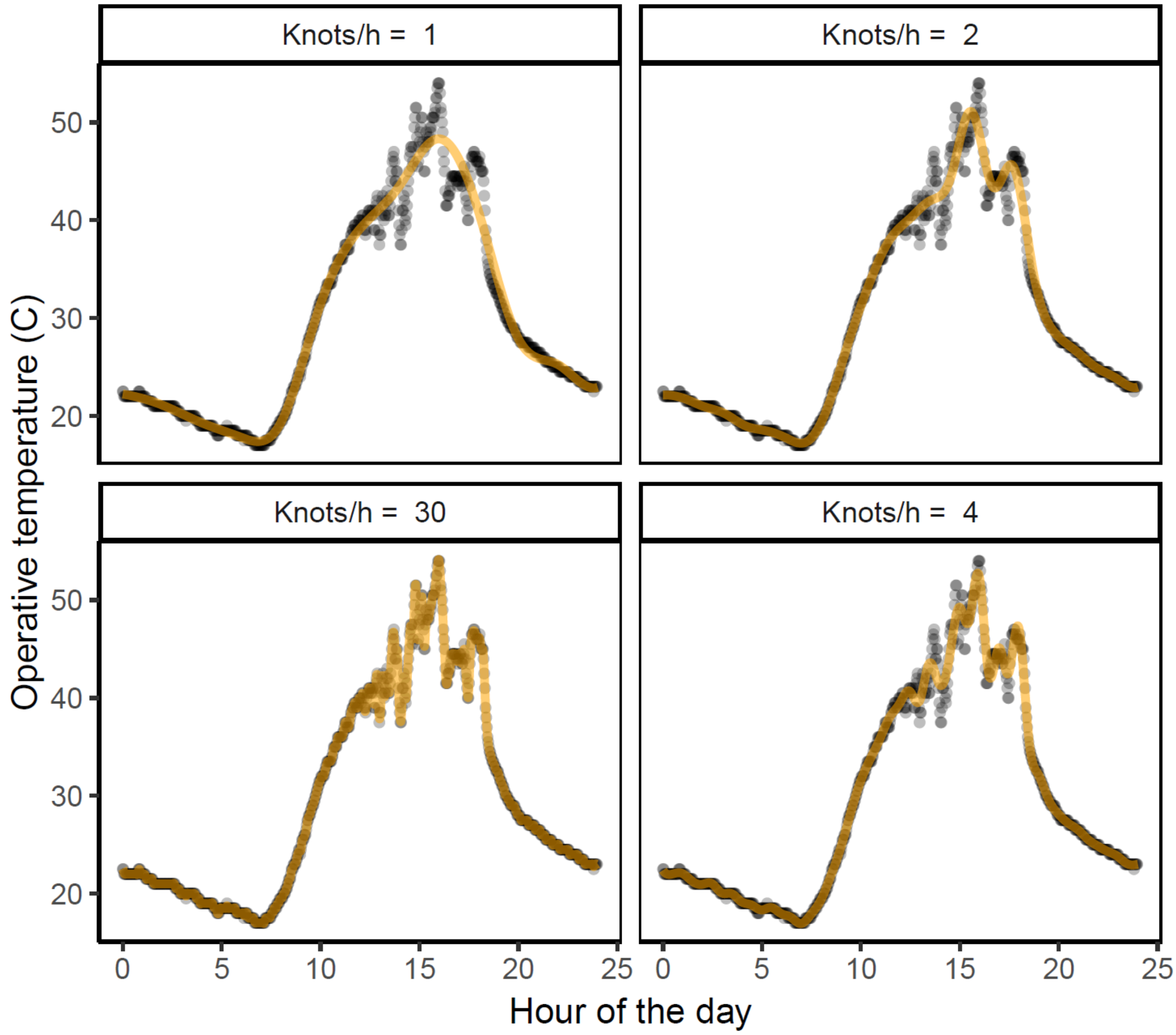

**Figure S14.** Example of a cubic spline (orange) fitted to OTM measurements (black) that logged every 2 minutes across 24 hours using 1, 2, 4 and 30 knots per hour.

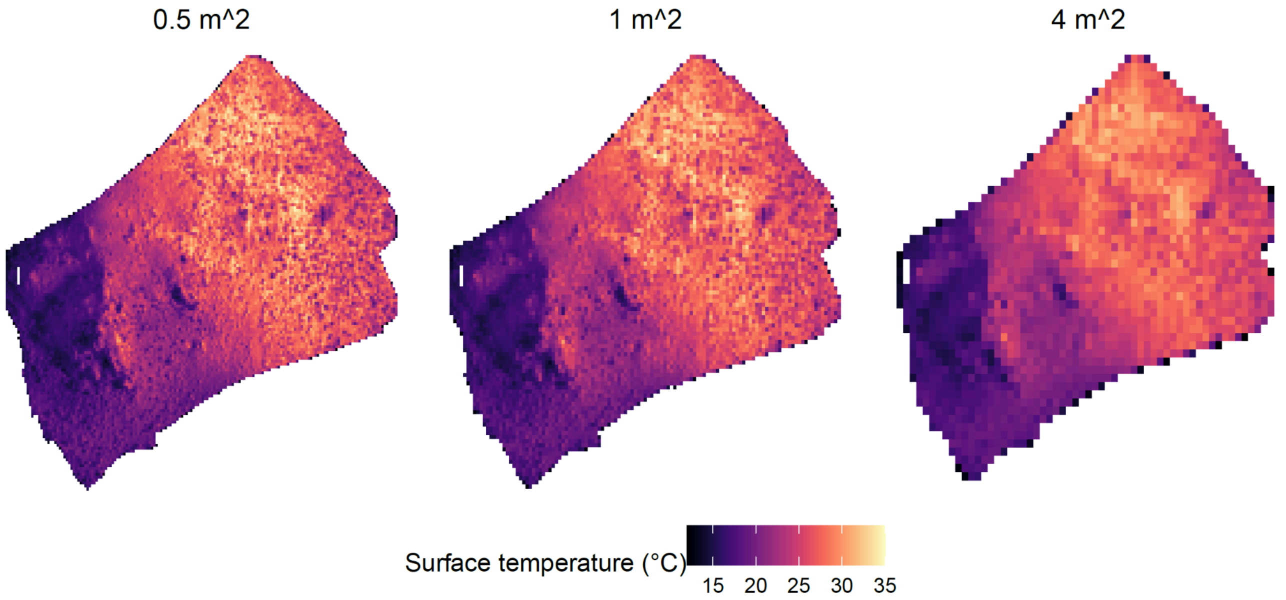

**Figure S15.** Results of processing the same flight at a spatial resolution of 0.5, 1 and 4 m^2.
